# RNA folding drives species-dependent host mRNA partitioning in Ebola virus condensates

**DOI:** 10.1101/2025.09.13.676059

**Authors:** Jingru Fang, Dylan C. Lam, Clarence K. Mah, Reika Watanabe, Hailey M. Tanner, Troy Comi, Yuan Cai, Cheng-Yi Chen, Aartjan J. W. te Velthuis, Clifford P. Brangwynne, Gene W. Yeo, Erica Ollmann Saphire

## Abstract

RNA viruses form membraneless condensates in host cells to drive replication, but whether these compartments regulate host RNAs remains unclear. Using MERFISH-based subcellular transcriptomics, we quantified cellular mRNA recruitment into Ebola virus condensates under basal and IFN-stimulated states. We find in the basal state, cellular RNAs with minimally folded coding regions are selectively recruited. Under IFN-stimulation, however, interferon-stimulated genes (ISGs) with structured 3′UTRs concentrate in viral condensates. We find both RNA structural features are conserved in the viral genome and support its retention in condensates. In parallel, for cellular mRNAs, we find partitioning into condensates prolongs RNA half-life and amplifies, rather than dampening, ISG expression. Fruit bats, which do not experience severe disease from RNA viruses, instead have ISGs with reduced 3′UTR folding, and may evade condensate-sequestration, enabling balanced antiviral responses. This selective stabilization links condensate function to RNA regulation as a molecular determinant of viral-host co-evolution and disease pathogenesis.

**In Brief:** Viral condensates are increasingly recognized as hubs for viral replication, but how they interact with host RNAs is poorly understood. This study reveals that Ebola virus condensates selectively recruit and stabilize host mRNAs, reshaping antiviral gene expression in ways that differ between humans and bats.

## Introduction

Cells organize biochemical reactions through dynamic, membrane-less compartments formed via phase separation. These condensates spatially coordinate essential processes such as ribosome biogenesis, RNA metabolism, and stress responses^1–3^. Many RNA viruses, including rabies, measles, and Ebola virus, have evolved to adopt or exploit this feature of our cells, forming viral condensates *de novo* within infected cells to compartmentalize genome replication and virion assembly^4–6^. In many RNA virus infections, the viral condensates are prominent cytoplasmic structures that can occupy up to 20–30% of the total cell volume^7–9^. While recent studies have begun dissecting the roles of viral condensates in viral genome replication and immune evasion^10,11^, it remains unknown whether and how these compartments influence host gene expression and infection outcome.

For almost all RNA viruses that form condensates, formation of these viral condensates is nucleated by viral RNA-binding proteins (RBP). These viral RBPs, such as nucleoproteins, oligomerize upon binding to the viral RNA genome during viral replication and packaging^6,11,12^. Multivalent interactions mediated by these viral RBPs combined with their high abundance during infection allow viral RBPs to be the scaffold of viral condensates. As infection progresses, the viral RBP concentration increases and reaches saturation that drives phase separation, creating a condensate phase enriched in scaffold proteins and viral RNA, surrounded by a dilute cytoplasmic phase^6^. These condensates serve as biochemical hubs, selectively recruiting and concentrating viral and host factors conducive for the completion of the viral life cycle.

RNA viruses that have a cytoplasmic replication cycle are thought to have limited access to the host transcriptional machinery, yet they can modulate host gene expression through multifunctional proteins that traffic to the nucleus^13–16^. Ebola virus protein VP24, for instance, antagonizes host innate immunity by interfering with the nuclear translocation of transcription factor STAT1 to block type 1 IFN signaling^17^. Paradoxically, transcriptomic analyses of viral infections often reveal robust and sustained innate immune activation, contributing to immunopathology in diseases such as rabies, measles, influenza, and Ebola virus disease (EVD)^18–22^. In fatal EVD cases, interferon-stimulated gene (ISG) expression is markedly elevated compared to survivors, and even convalescent EVD patients exhibit persistent inflammatory gene signatures years after recovery^23,24^. This discrepancy between viral immune evasion strategies and host immunopathology suggests a missing mechanistic link between viral biology and regulation of host gene expression. Interestingly, bats—natural reservoirs for many RNA viruses including Ebola virus—can tolerate high viral loads without exhibiting severe disease, often mounting a controlled interferon response with limited inflammatory pathology^25–28^. How bats achieve this balance, while other mammals succumb to immune-driven tissue damage remains a focal point of current research. Understanding how viral condensates might differentially modulate host gene expression and immune responses across species could provide critical insight into virus-host co-adaptation and species-specific viral pathogenesis.

A defining feature of viral RBPs for negative-strand RNA viruses is their sequence-independent RNA-binding activity. For instance, Ebola virus nucleoprotein (NP) uniformly coats the single-stranded viral genome to form viral nucleocapsids^29^, while the viral polymerase cofactor VP35 engages double-stranded viral RNA structures to facilitate replication and evade innate immunity^30,31^. However, this lack of sequence specificity enables viral RBPs to also bind cellular mRNAs, which far outnumber viral RNAs in infected cells, particularly early in infection. Biochemical, imaging, and structural studies have detected cellular RNAs within viral condensates and bound by purified viral RBPs^32–34^, but the specific identities of these mRNAs, their relative enrichment in viral condensates, and the functional consequences of the binding of these RNAs by viral RBPs during infection remain unknown. Whether viral condensates passively sequester host mRNAs or actively reshape the host transcriptome through selective partitioning is an open question with significant implications for virus-host interactions.

Here, we show the sequence-independent RNA-binding activity of viral RBPs underlies the aberrant partitioning of host mRNAs into viral condensates, thereby altering their subcellular distribution and function in gene expression. Using Ebola virus as a model system, we investigated how viral condensates engage host transcripts under basal and type I interferon-stimulated conditions and compared differential partitioning in human and bat cells. We applied multiplexed error-robust fluorescence *in situ* hybridization (MERFISH) to spatially resolve and quantify host mRNAs at single-cell resolution, enabling a mechanistic dissection of their partitioning behavior. Our findings uncover how viral condensates of negative-strand RNA viruses sequester host mRNAs through a mechanism shaped by both RNA folding and host IFN responses, revealing an unconventional layer of virus–host interactions that link condensate biology to viral pathogenesis and immune dysregulation.

## Results

### Ebola Viral Condensates Partition Host mRNAs in Human and Bat Cells

We asked whether Ebola viral condensates, reconstituted by co-expressing two scaffold proteins, NP and VP35, can recruit cellular mRNAs. Using poly(A)-FISH to detect mRNAs, we visualized viral condensates via mNeonGreen-tagged VP35 (mNG-VP35), which we previously validated as a functional marker for Ebola viral condensates^35^. Immunofluorescence staining confirmed co-localization of NP within these condensates. However, due to antibody accessibility differences, NP immunofluorescence staining was unequal across the condensate (stronger in the periphery; weaker in the interior; Figure 1A). Thus, we used the mNG-VP35 signal, which was homogeneous across each condensate, to map viral condensates.

**Figure 1.**
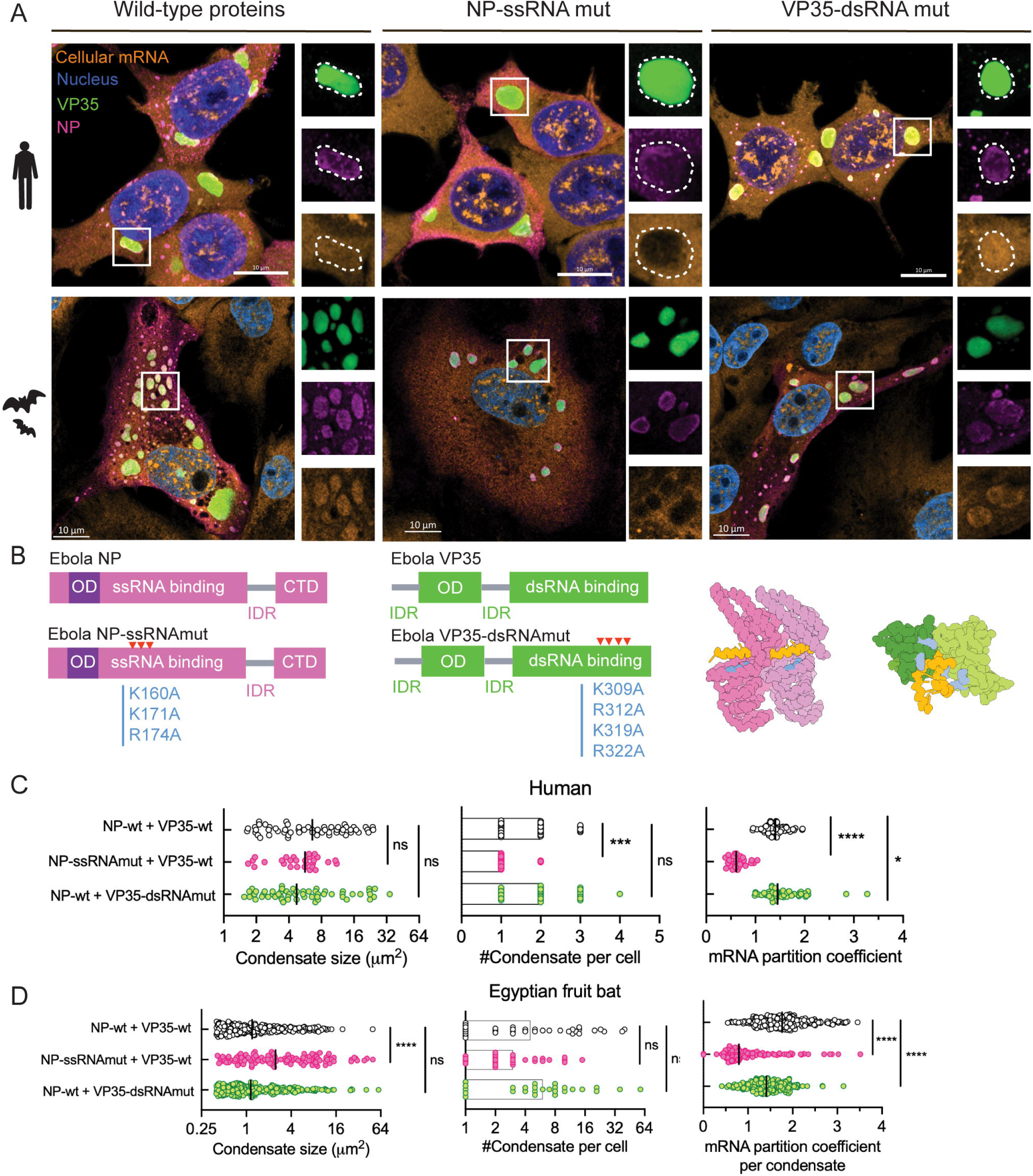
Ebola virus condensates partition cellular mRNA in human and fruit bat cells. **A.** Confocal microscopy of fixed and stained HEK293T (human kidney) and RAKSM (bat kidney) cells co-expressing untagged Ebola virus NP and mNG-tagged Ebola virus VP35 in either wild type or RNA-binding mutant. Individual channels are shown for a representative condensate in each condition. Scale bar: 10 μm. Nuclei stained with Hoechst. **B.** Schematic of Ebola NP and VP35 wild type or RNA-binding mutants with amino acid changes marked in RNA-binding domain and colored in blue in existing NP (model in pink, PDB:8USN^47^) and VP35 structure (model in green, PDB: 3L25^30^). OD: oligomerization domain; IDR: intrinsically disordered region; NP/VP35-bound RNA shown in yellow. **C. D.** Condensate size, number, and cellular mRNA partition coefficient for Ebola virus NP-VP35 condensates with either wild type or RNA-binding mutant in human or bat cells. Median values with individual data points (per condensate) from two experiments (> 12 cells per condition per experiment) are shown. ****, P<0.0001; ***, P<0.001, *, P<0.05, ns: not significant, by Mann-Whitney test.

In HEK293T cells, poly(A)-FISH revealed robust mRNA signals within NP–VP35 condensates, suggesting cellular mRNAs co-localize in these viral compartments. To determine whether this mRNA recruitment depends on the RNA-binding activity of NP and VP35, we transfected cells with well-characterized RNA-binding mutants that ablate NP-ssRNA^36,37^ or VP35-dsRNA interactions^38,39^, respectively (Figure 1B). Each mutant retained condensation ability, though differentially, when co-expressed with the wild-type scaffold partner (Figure 1C).

Condensates containing the NP-ssRNA binding mutant failed to retain cellular mRNAs. In contrast, condensates containing the VP35-dsRNA binding mutant retained cellular mRNAs, exhibiting mRNA partitioning comparable to wild-type levels (Figure 1A). Quantification of the mRNA partition coefficient, defined as the ratio of mRNA-FISH intensity in condensates versus cytoplasm, shows a median of 1.4 for wild-type condensates. This drops to 0.61 (42% of wild-type, P<0.0001, Mann-Whitney test) in condensates with NP-ssRNA mutant plus wild-type VP35, but increased to 1.45 (103% of wild-type, P=0.0185, Mann-Whitney test) in condensates containing wild-type NP plus the VP35-dsRNA mutant (Figure 1C). These results indicate NP’s ssRNA-binding activity primarily drives cellular mRNA recruitment into Ebola viral condensates.

Next, we examined whether this mRNA partitioning phenotype is conserved in cells from viral reservoir host. Given the absence of a definitive reservoir host for Ebola virus, we utilized immortalized kidney cells (RAKSM) from Egyptian fruit bats (*Rousettus aegyptiacus*), a species that serves as the natural reservoir for the closely related Marburg virus^40–42^. Similar to human cells, NP-VP35 co-expression also formed cytoplasmic condensates in bat cells with varying numbers and sizes (Figure 1D).

In bat cells, however, wild-type condensates exhibited a higher mRNA partition coefficient (median = 1.76) compared to human cells (median =1.4), implying these bat cells recruit more cellular mRNA to condensates than do human cells. In the bat cells, disruption of NP ssRNA-binding reduced mRNA partitioning to 0.79 (45% of wild-type, P<0.0001, Mann-Whitney test) similar to observations in human cells. However, in the bat cells, VP35-dsRNA binding mutant condensates showed a minor decrease to 1.41 (80% of wild-type, P<0.0001, Mann-Whitney test) (Figure 1D). This observation contrasts with that of human cells in which the VP35-dsRNA-binding mutant had no effect on mRNA partitioning. Thus, in human cells, NP dominates mRNA recruitment, whereas in the bat cells, both NP and VP35 contribute to polyadenylated mRNA partitioning. Together, these results suggest some host-specific differences in the interactions between viral condensates and cellular mRNAs.

To test whether our observations were specific to the cell line used, we also performed poly(A)-FISH analysis in human hepatocyte Huh7 cells, relevant to Ebola virus tropism, and VeroE6 African green monkey kidney cells, commonly used for viral propagation (Figure S1A). In Huh7 cells, the global mRNA partitioning pattern in viral condensates closely mirrored that in HEK293T cells: the median partition coefficient was similar in wild-type condensates (1.46), and the NP ssRNA-binding remained the primary determinant of cellular mRNA recruitment (Figure S1B). In VeroE6 cells, global mRNA partitioning was modestly reduced (median = 1.24), but NP ssRNA binding remained determinant of mRNA recruitment (Figure S1C).

Having established that cellular mRNAs are constituents of viral condensates, we next investigated their functional roles in viral condensate behavior. Previous studies have shown that cellular RNA promotes phase separation for endogenous condensates, like stress granules^43^, in uninfected cells^44^. Hence, we proposed that binding of cellular mRNAs by viral proteins could enhance viral condensate stability. These interactions may allow viruses to maintain replication-competent compartments, especially at the onset of infection when viral RNA copies are low.

To test this hypothesis, we exposed condensates—with or without cellular mRNAs, using either wild-type or mutant NP—to hypotonic shock, a transient perturbation achieved by diluting intracellular proteins and reducing macromolecular crowding with water to dissolve liquid-like condensates in live cells^45,46^. We quantified condensate dissolution by measuring the coefficient of variation (CV) of VP35 fluorescence intensity across cells. Wild-type condensates, which retain cellular mRNAs, resisted dissolution following hypotonic shock (by diluting media 2 times in water, ∼110 mOsm), although partial disassembly was observed. In contrast, condensates formed by the NP RNA-binding mutant, lacking mRNA recruitment, disassembled to a greater extent (Figure S2A).

Live-cell imaging further revealed that condensates with wild-type NP dissolved gradually over several minutes post-osmotic shock (Video S1), while condensates with mutant NP disintegrated rapidly within seconds (Video S2 and Figure S2B). These findings suggest that cellular mRNA binding enhances the physical resilience of Ebola viral condensates under osmotic stress, potentially stabilizing viral replication compartments during infection.

### Ebola Virus Condensates Enrich Single-Stranded Cellular mRNAs with Minimal Coding Sequence Structure at Steady State

We next asked whether Ebola virus condensates selectively enrich specific cellular mRNAs. Using MERFISH^48^, we quantified transcript abundance and subcellular localization in HEK293T cells with reconstituted Ebola NP-VP35 condensates. MERFISH allows amplification-free quantification of thousands of RNA species simultaneously at single-cell resolution, providing multidimensional data to assess RNA partitioning in condensates. RNA partitioning is represented by the transcript density ratio, calculated by normalizing transcript density (normalized to area) in viral condensates to that in the cytoplasm. A transcript density ratio greater than 1 indicates mRNA enrichment in condensates (Figure 2A).

**Figure 2.**
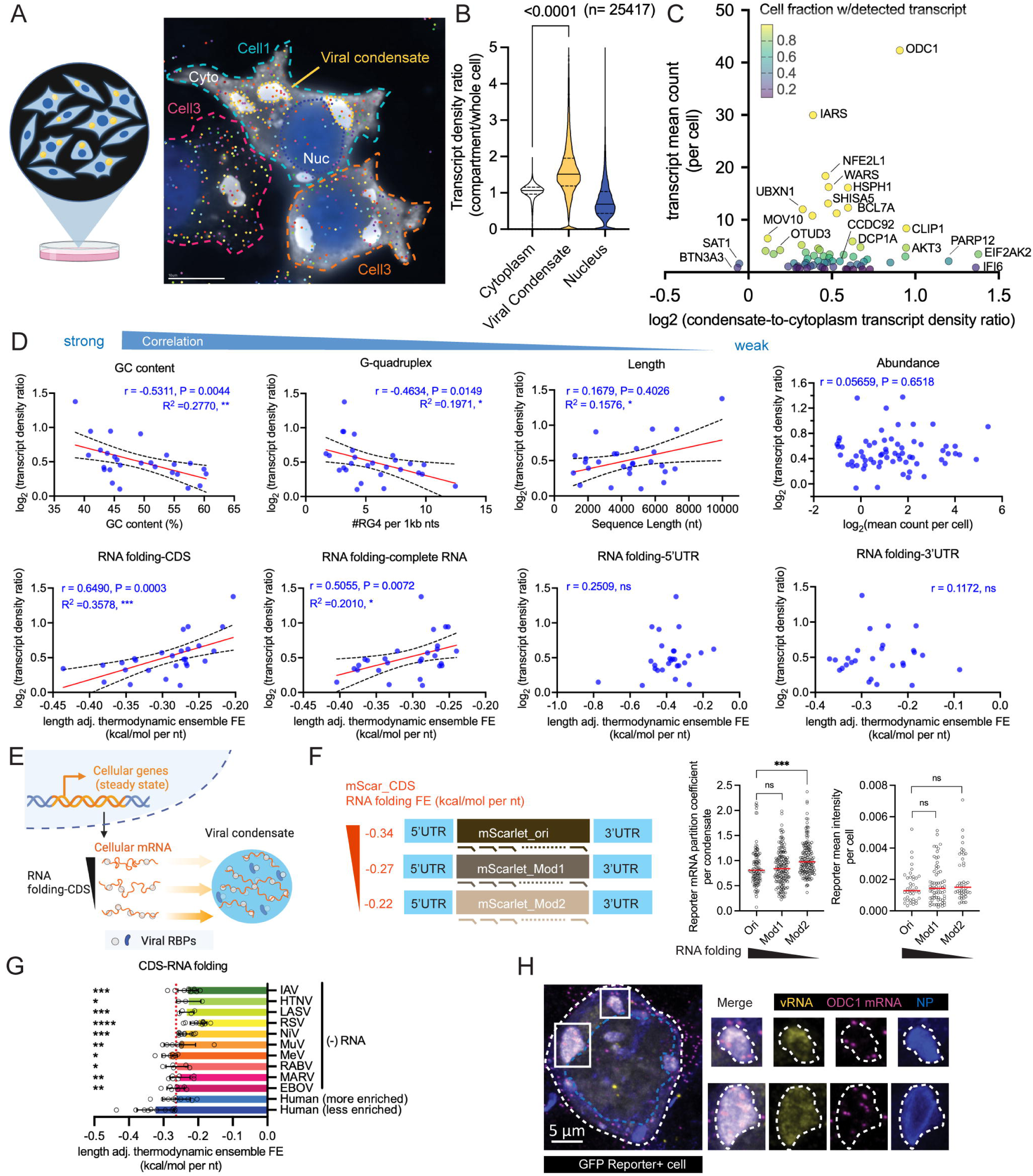
Cellular mRNAs are differentially enriched in Ebola virus condensates See also in Figure S3, Table S1, Table S4. **A.** Schematic of HEK cells expressing Ebola virus condensates used in the MERFISH experiment. Individual cells, cytoplasm, viral condensates, and nucleus are annotated in a representative field of view in MERFISH raw data, with detected transcripts shown in different colors. Scale bar: 10 µm. **B.** Comparison of MERFISH-detected, transcript density ratio (compartment-to-whole cell) in different subcellular compartments for 25,417 cells analyzed. Solid line: median. Dashed line: quartiles. Paired t-test used to determine difference between cytoplasm and viral condensate with P value indicated on top. **C.** Distribution of log_2_transcript density ratio (condensate-to-cytoplasm) over abundance for top 66 high-confidence transcripts detected by MERFISH. Percentage of cells with detected transcripts scaled by color map. **D.** Correlation analysis between RNA features and condensate enrichment depicted by transcript density ratio. Both Spearman r and P values from Spearman correlation analysis are shown for all conditions. ns, not significant. R^2^, P values (*, P<0.05, **, P<0.01, ***, P<0.001) and fitted lines with 95% confidence intervals from linear regression are shown for significant correlations. **E.** Schematic illustration of selective transcript enrichment in viral condensates for transcripts with minimal RNA folding in CDS, under steady state conditions. RBPs: RNA-binding proteins. Created using BioRender. **F.** Schematics of synthetic RNA reporters composed of the same 5’UTR and 3’UTR of human ODC1 mRNA and different mScarlet coding sequences with varying level of predicted length adjusted RNA folding free energy as annotated in the unit of kcal/mol per nt on the side. Different sets of SABER-FISH probes are designed to label each mScarlet coding sequence variant. Representative results of two experiments (>39 cells in each condition in total) are shown. Mann-Whitney test used to determine the difference in either reporter mRNA partition coefficient in viral condensate or mScarlet reporter mean intensity between mScarlet reporter carrying the original and modified coding sequences. The annotated mScarlet reporter sequence is shown in Table S4. **G.** Comparison of the predicted RNA-folding energy (length-adjusted) for cellular and negative-strand virus CDS regions. Human (more/less enriched): high-confidence cellular mRNAs detected in MERFISH with top/bottom 50% ranked transcript enrichment in the viral condensates. HTNV: Andes hantavirus, LASV: Lassa virus, IAV: Influenza A virus, RSV: Respiratory syncytial virus, MARV: Marburg virus, MuV: Mumps virus, NiV: Nipah virus, RABV: Rabies virus, MeV: Measles virus, EBOV: Ebola virus. Welch’s t test used to compare the difference in RNA folding of human (less enriched) cellular mRNAs versus each viral RNA (negative sense). **H.** Confocal microscopy of fixed and stained HEK293T cell transfected with Ebola virus minigenome system and active for viral RNA synthesis indicated by fluorescence reporter. Individual channels of magnified viral condensates are shown for reporter viral RNA and endogenous ODC1 mRNA stained by SABER-FISH, viral condensates marked by immunofluorescence-stained Ebola virus NP protein. A representative cell from two experiments (>7 reporter+ cells per experiment) is shown. Scale bar: 5 µm in merge overview image, 2 µm in magnified images.

We designed MERFISH probes targeting 136 disease-relevant genes to test our hypothesis Ebola virus can differentially partition host mRNAs into viral condensates, and this partitioning could assist post-transcriptional regulation of host gene expression. The 136 genes include Ebola-specific IFN-stimulated genes (ISGs)^49^, differentially expressed genes from Ebola virus infection *in vivo*^50^, and conserved mammalian innate immune genes^51^.

We analyzed ∼90,000 cells and found ∼30% contained detectable viral condensates. Viral condensates were enriched in cellular mRNAs encoded by the 136-gene panel, while nuclear regions were depleted of these cellular mRNAs (Figure 2B), suggesting these mRNAs were sampled at their steady state post-transcription. The magnitude of enrichment of individual mRNAs was consistent with mRNA-FISH measurements of bulk mRNA, though MERFISH revealed greater heterogeneity among individual mRNAs (Figure 2B), compared with the ensemble of mRNA measured by poly(A)-FISH (Figure 1C), suggesting transcript-specific partitioning.

To map transcript-specific partitioning in viral condensates, we ranked all MERFISH-detected transcripts by mean-counts per cell and selected 66 high-confidence hits with mean-counts per cell higher than that of misidentification controls (Figure S3A). Since most ISGs are only expressed upon IFN treatment, they have a minimal representation in the 66 high-confidence transcripts for the non-IFN-stimulated experimental setting. Among the 66 high-confidence transcripts, only SAT1 and BTN3A3 are not enriched in condensates. The remaining 64 transcripts exhibit varying degrees of abundance (mean count = 0.5 to 40 per cell) and enrichment in viral condensates (Figure 2C). Notably, we do not observe a correlation between mRNA abundance and enrichment in viral condensates, suggesting mRNA partitioning can be selective rather than mediated by concentration-dependent passive diffusion (Figure 2D).

To independently validate MERFISH-measured RNA partitioning, we performed RNA co-immunoprecipitation (RNA-IP) using whole-cell lysates from HEK 293T cells co-expressing the same viral scaffolds (mNG-VP35 and NP) used in MERFISH experiments. We quantified transcript enrichment in condensates by RT-qPCR for five representative genes (SAT1, MOV10, IARS, ODC1, EIF2AK2). The RNA-IP enrichment strongly correlates with the MERFISH-derived transcript density ratio (R² = 0.92, P = 0.0098) (Figure S3B). We further confirmed condensate recruitment of ODC1 mRNA via SABER (Signal Amplification By Exchange Reaction)-FISH^52^, which is ablated by disrupting NP ssRNA-binding activity (Figure S3C).

To uncover molecular features governing RNA partitioning, we analyzed seven RNA parameters across a subset of 27 condensate-enriched, high-confidence transcripts, defined by mean counts per cell >3 and condensate-to-cytoplasm transcript density ratio >1 (log_2_transcript density ratio >0) (Figure 2D). We found the GC content and the number of predicted G-quadruplex motifs negatively correlated with condensate partition, whereas RNA length weakly but positively correlated with condensate partition, gated by the limited variation in length for selected transcripts. In line with previously observed RNA structure biases, RNAs with higher single-strandedness, measured by length-normalized free energy predicted for RNA folding ensemble^53–55^, positively correlated with condensate partition. We normalized the free energy for RNA folding by RNA length to minimize the contribution of length alone in condensate partition. Partitioning correlated most significantly with RNA folding in the coding sequences (CDS), rather than untranslated regions (UTRs), suggesting that single-stranded regions within the CDS interact with the viral-ssRNA binding protein NP and drive condensate localization (Figure 2E).

Next, we tested whether CDS folding alone is sufficient to drive mRNA partitioning in viral condensates using synthetic reporter mRNAs. To minimize chemical differences associated with in-vitro transcribed mRNAs, which lack most endogenous RNA modifications, we generated reporter transcripts by plasmid-based expression driven by a strong Pol II promoter and co-expressed each reporter with the viral condensate scaffold proteins (NP and VP35). Reporters contain the 5’ and 3’ UTRs of ODC1, an endogenous transcript highly enriched in viral condensates, flanking mScarlet CDS.

The native mScarlet CDS is predicted to be highly structured (length-normalized folding energy = −0.34 kcal/mol/nt). To test whether CDS folding influences condensate recruitment, we generated two synonymous mScarlet variants with reduced predicted secondary structure (−0.27 and −0.21 kcal/mol/nt) by replacing selected GC-rich codons with AT-rich codons. Reporter localization was quantified by RNA-FISH using probes targeting each mScarlet CDS, and condensate enrichment was measured by partition coefficient (Figure 2F).

If reduced RNA folding promotes condensate recruitment, the least structured transcript should exhibit the highest partition coefficient. Consistent with this prediction, RNA-FISH revealed progressively increased partitioning of less-structured mScarlet reporter mRNAs into viral condensates (Figure 2F). Importantly, all variants showed comparable protein expression based on mean mScarlet fluorescence, indicating differences in condensate partitioning were unlikely to arise from altered translation or polysome occupancy.

These reporters have limitations: the mScarlet CDS (696 nt) is substantially shorter than endogenous transcripts enriched in viral condensates (average length = 1,896 nt), likely reducing the number of potential NP interactions. This reduced valency may explain their relatively modest partitioning. To test the robustness of the correlation between CDS folding and condensate partitioning, we designed a locked nucleic acid (LNA)/DNA mixer antisense oligo targeting a predicted single-stranded region of endogenous ODC1 mRNA. LNA binding is resistant to RNase-H cleavage and will occlude viral NP ssRNA-binding. Increasing doses of ODC1-targeting LNA reduces endogenous ODC1 partitioning in viral condensate without significantly affecting total ODC1 mRNA levels (Figure S3D). Together with the reporter assays, these results support our conclusion that reduced CDS secondary structure promotes RNA partitioning into viral condensates.

Given that one role of viral condensates is to act as a hub for negative-strand RNA virus replication, we hypothesized the viral genome sequence would contain minimal secondary structure to optimize condensate retention. To test this hypothesis, we analyzed the CDS of 10 negative-strand RNA viruses, Andes Hantavirus, Lassa, Influenza A, Respiratory syncytial virus, Marburg, Mumps, Nipah, Rabies, Measles, and Ebola viruses. We indeed found that the CDS regions of all 10 viruses exhibit a lower degree of secondary structure than do MERFISH-detected cellular mRNAs, which have low condensate partitioning (Figure 2G, Figure S3E, all viral RNAs compared to Human-less enriched). Negative-strand RNA viruses encode their genome in the negative orientation, but also produce complementary, non-coding, anti-genome, positive-strand RNA in replication, and capped, positive-strand mRNAs in transcription. Interestingly, the CDS folding of the negative sense genomes more closely resembles condensate-enriched cellular mRNA than do the positive sense anti-genome/mRNA; there may be more selection pressure to drive and retain the genome into the condensates (Figure 2G and Figure S3E). Together, these results imply thermodynamic constraints that control the RNA composition in Ebola viral condensates are applicable to both host and viral genomic RNAs.

To confirm cellular mRNA recruitment in viral condensates in the presence of viral RNA, we assessed the localization of a representative cellular transcript, ODC1, in viral condensates during active viral RNA synthesis. Using a replication-competent Ebola minigenome reporter system^35,56^, we detected ODC1 mRNA alongside viral genomic RNA (vRNA) within condensates of reporter-positive HEK293T cells (Figure 2H). In this system, the GFP reporter signal is produced by the negative-sense viral minigenome, replicated and transcribed by the co-expressed viral polymerase L and translated by host ribosomes (Figure S3F). This experiment indicates condensate-mediated recruitment of cellular mRNAs can occur even during active viral RNA synthesis. Together, our data reveal that Ebola virus condensates selectively enrich cellular mRNAs with minimal CDS folding, a property that aligns with the structural features of the viral genome itself.

### Upon type 1 IFN stimulation, Ebola virus condensates enrich mostly cellular mRNAs encoded by IFN-stimulated genes (ISGs)

While dysregulation of immune responses is a hallmark of fatal Ebola virus disease^24^, the molecular basis for how Ebola virus perturbs host immune homeostasis remains unclear. Having established the presence of cellular transcripts in Ebola viral condensates, we hypothesized these condensates aberrantly partition ISG mRNAs, thereby modulating their fate during infection, and contributing to immune dysregulation. To test this hypothesis, we quantified the cellular mRNA partitioning landscape in the context of a type I IFN response^57^, which broadly activates antiviral programs in non-immune cells, including the immortalized kidney cells used in this study.

We first performed poly(A)-FISH to quantify global mRNA partitioning in Ebola viral condensates in HEK cells stimulated with type I IFN (Figure 3A). We confirmed IFN stimulation by detecting STAT1 phosphorylation and induction of ISG15 and RIG-I, both known ISGs in human and Egyptian fruit bats (Figure S4A). In IFN-stimulated human cells, the median mRNA partition coefficient for wild-type viral condensates was 1.38. This coefficient dropped to 0.71 (52% of wild-type, P<0.0001, Mann-Whitney test) in condensates formed by NP-ssRNA mutant plus wild-type VP35, but increased to 1.68 (122% of wild type, P<0.0001, Mann-Whitney test) in condensates containing wild-type NP and VP35-dsRNA mutant (Figure 3B). In IFN-stimulated cells, NP that cannot bind mRNA lost its ability to enrich cellular mRNAs in the viral condensates. In contrast, VP35 that cannot bind dsRNA exhibits increased mRNA partitioning into the condensates. Hence, NP ssRNA-binding activity remains important for RNA partitioning in interferon-stimulated cells, while VP35 dsRNA-binding activity limits RNA partitioning in interferon-stimulated cells.

**Figure 3.**
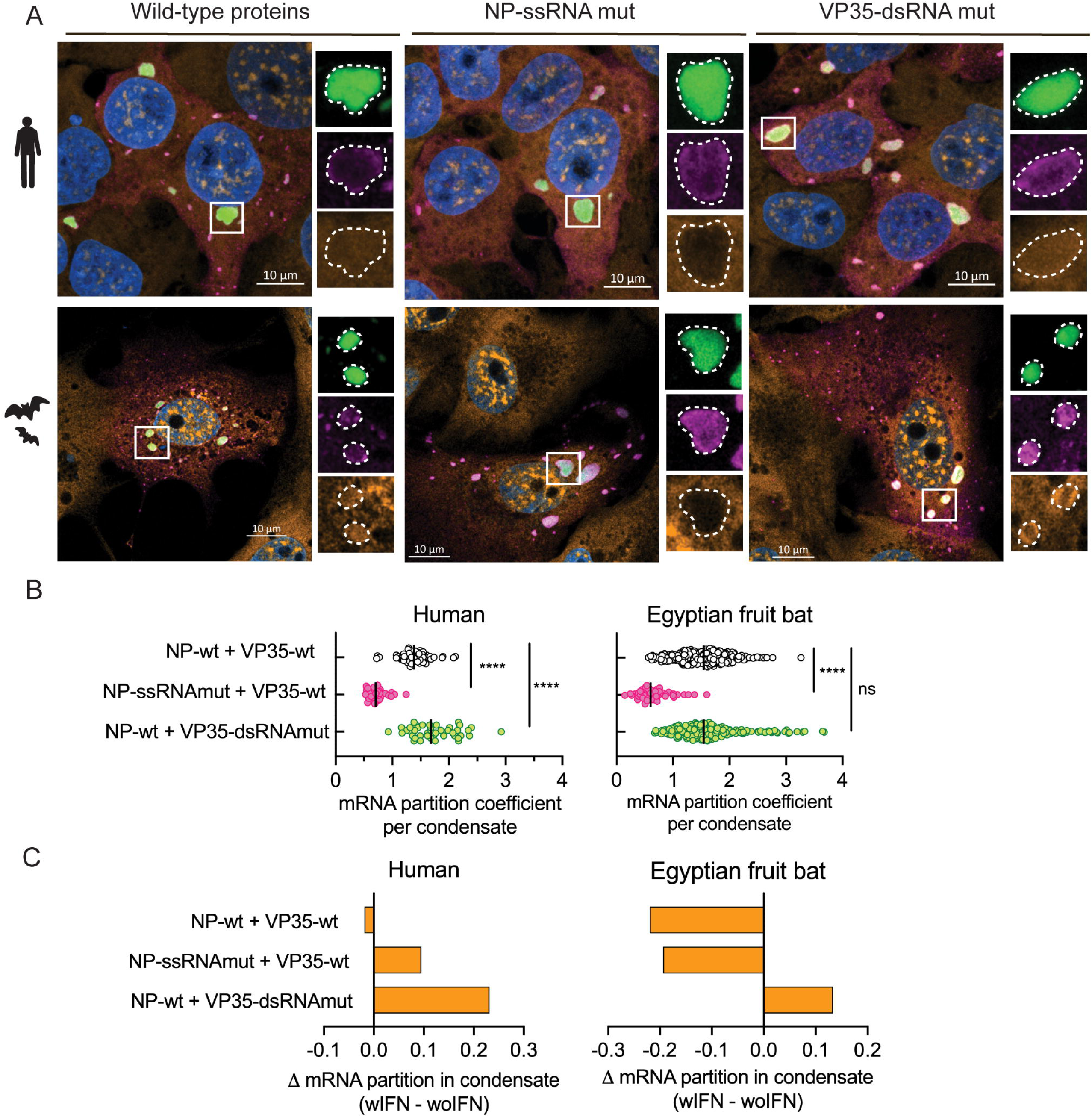
Ebola virus condensates partition cellular mRNAs in human and fruit bat cells under type 1 IFN stimulation. **A.** Confocal microscopy of fixed and stained HEK293T (human kidney) and RAKSM (bat kidney) cells co-expressing untagged Ebola virus NP and mNG-tagged Ebola virus VP35, in either wild-type or RNA-binding mutant formats, and subsequently incubated with type 1 IFN (1000U/ml) for 24 hours. Individual channels are shown for a representative condensate in each condition. Nuclei stained with Hoechst. Scale bar: 10 µm. **B.** Cellular mRNA partition coefficient in Ebola virus NP-VP35 condensates, with either wild-type or RNA-binding mutant scaffolds, in IFN-stimulated human or bat cells. Median values with individual data points (per condensate) from two experiments (with > 7 cells in each condition per experiment) are shown. ****, P<0.0001; ns, not significant, by Mann-Whitney test. **C.** Changes in cellular mRNA partitioning in viral condensates in IFN-stimulated cells compared to unstimulated cells. Difference in median partition coefficients for each condition is shown.

We next compared mRNA partitioning in bat cells and human cells and observed similar trends. In type I IFN-stimulated Egyptian fruit bat cells, wild-type NP plus wild-type VP35 exhibits a median mRNA partition coefficient of 1.54 and the NP ssRNA-binding mutant exhibits an mRNA partition of 0.6 (39% of wild-type, P<0.0001, Mann-Whitney test). However, mRNA partitioning in condensates with the VP35-dsRNA mutant was identical to the wild-type control, suggesting VP35-dsRNA binding activity is no longer important for binding cellular RNA under interferon stimulation (Figure 3A and 3B). We reasoned that cellular RBPs residing in viral condensates may replace or compete with Ebola VP35 for dsRNA binding, given that multiple cellular dsRNA-binding proteins (e.g., PKR, STAU1, ADAR) have been found to interact with viral proteins inside Ebola viral condensates^58–60^. IFN stimulation likely enhances the expression, condensate recruitment, or RNA-binding activity of these cellular RBPs^61^, and shifts the cellular mRNA composition of viral condensates.

In human cells, following IFN stimulation, RNA partitioning became less dependent on NP-mediated RNA binding activity, supporting alternative binding forces mediated by cellular RBPs. However, in bat cells, this dependency is flipped, with a higher contribution of NP-mediated RNA binding in IFN-induced cellular mRNA partition than in steady state (Figure 3C).

In addition to the bulk quantity of mRNA partitioned into condensates, we observed clear spatial differences in how RNA partitions in viral condensates formed in human vs. bat cells. In human cells, cellular mRNAs are homogeneously distributed in condensates formed by both wild-type and all NP and VP35 mutants. In bat cells, while cellular mRNAs are homogeneously distributed with wild-type VP35, they are unevenly enriched in the periphery of viral condensates containing the VP35 mutant (Figure 3A).

Next, to map transcript-specific rather than overall partitioning in viral condensates, we performed a MERFISH-based subcellular analysis in type 1 IFN-stimulated human cells reconstituted with Ebola virus NP-VP35 condensates. We analyzed transcripts encoded by the same 136 disease-relevant genes described above. Similar to bulk mRNA measurements, MERFISH analysis reveals a modest enrichment of transcripts overall in condensates (median = 1.13) relative to the cytoplasm (median = 1) (Figure 4A). Despite a smaller sample size (n = 2419 cells), we recovered 81 high-confidence transcripts (mean count greater than misidentification controls) in the 136-disease-relevant gene panel (Figure 4B). These high-confidence transcripts include over 20 IFN-stimulated gene (ISG) transcripts that were undetectable in the absence of type 1 IFN stimulation.

**Figure 4.**
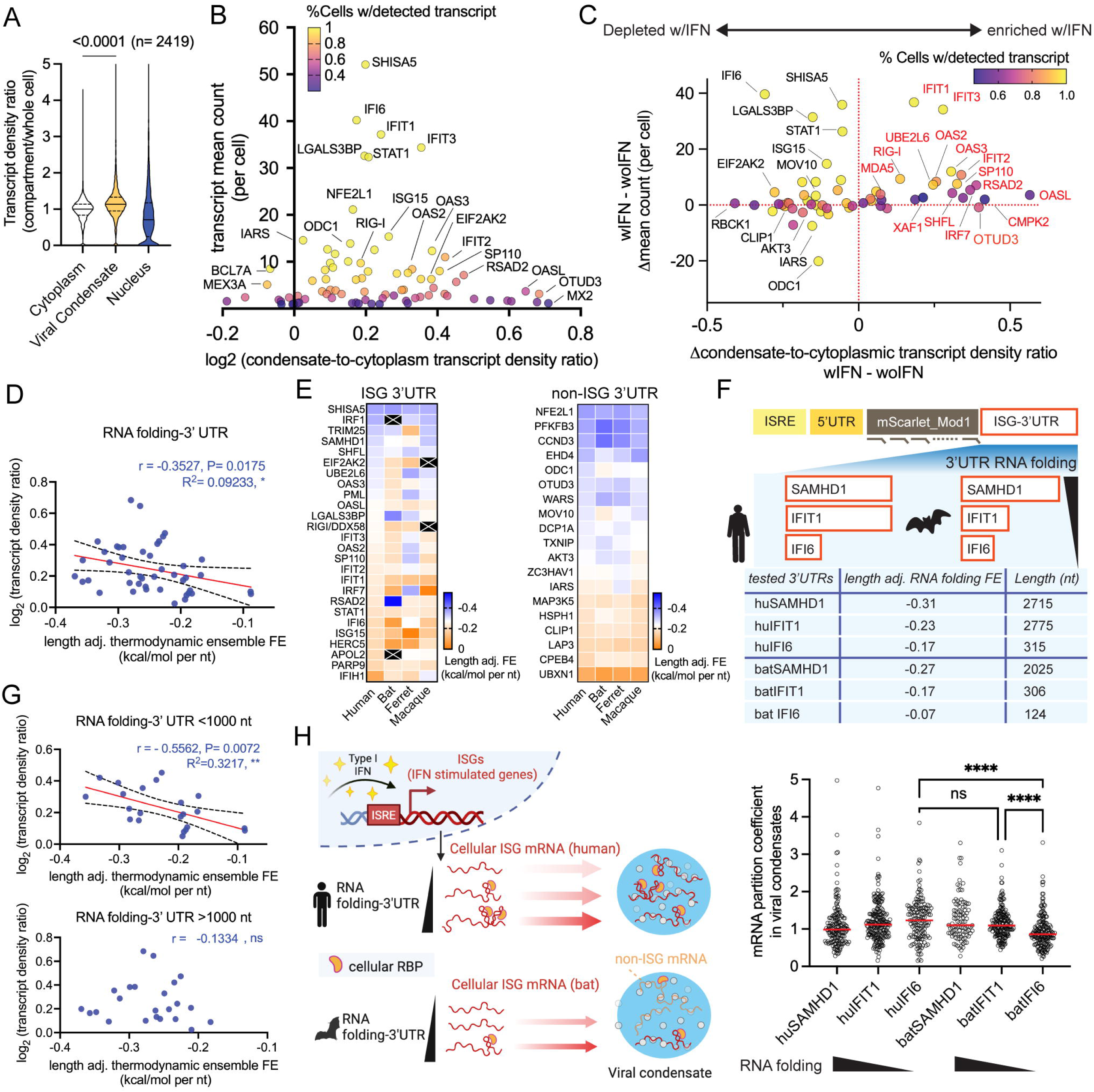
Interferon-stimulated gene (ISG) encoded mRNA partitioning in Ebola virus condensates upon type 1 IFN stimulation See also in Figure S4, Table S1, Table S4. **A.** Comparison of MERFISH-detected transcript density ratio in different subcellular compartments for 2,419 IFN-stimulated cells analyzed. Solid line: median. Dashed line: quartiles. Paired t-test used to determine difference between cytoplasm and viral condensate with P value indicated on top. **B.** Distribution of log_2_transcript density ratio (condensate-to-cytoplasm) over abundance for top 81 high-confidence transcripts in IFN-stimulated cells detected by MERFISH. Percentage of cells with detected transcripts scaled by color map. **C.** IFN-induced changes in MERFISH-detected transcript density ratios and transcript mean count per cell are shown for the top 60 transcripts detected by MERFISH with IFN-stimulated cells. Transcripts labeled in red are undetectable without IFN-stimulation and have their transcript density ratios from steady state assigned as 0. **D.** Correlation analysis between the length-adjusted 3’UTR RNA folding energy (predicted) and condensate enrichment depicted by transcript density ratio. Spearman r and P values from spearman correlation analysis are shown. R^2^, P values (*, P<0.05) and fitted lines with 95% confidence intervals from linear regression are shown. **E.** Comparison of the predicted 3’UTR RNA folding among human, bat, ferret, and macaque orthologs for 26 ISG and 19 non-ISG transcripts within 45 condensate-enriched hits. ISG: defined by > 2 fold change in transcript mean count, from IFN-stimulated to unstimulated samples, as measured in MERFISH subcellular analysis. Black: Genes that are not identified (IRF1, APOL2, RIGI) or with <10 nt 3’UTR (EIF2AK2) encoded by a given species. **I.** Synthetic RNA reporters composed of the same ISRE, 5’UTR and mScarlet-mod1 CDS with different 3’UTR encoded by human or bat ISG. For each 3’UTR sequence used, the corresponding level of predicted, length-adjusted, RNA-folding free energy is annotated as kcal/mol per nt. The same set of SABER-FISH probes targeting the mScarlet-mod1 CDS were used for FISH analysis. Representative results of two experiments (>39 cells in each condition in total) are shown. Mann-Whitney test used to determine the difference in reporter mRNA partition coefficient in viral condensate between reporters carrying selected 3’UTR sequences. The annotated reporter sequence is shown in Table S4. **F.** Correlation analysis between the length-adjusted 3’UTR RNA-folding energy (predicted) and condensate enrichment depicted by transcript density ratio for 3’UTRs that are shorter/longer than 1000 nucleotides. Spearman r and P values from spearman correlation analysis are shown. R^2^, P values (*, P<0.05) and fitted lines with 95% confidence intervals from linear regression are shown. **G.** Schematic illustration of selective partitioning of cellular mRNAs in viral condensates for RNAs with more structured 3’UTR (under type 1 IFN stimulation). Created using BioRender.

To compare transcript exchange in viral condensates between basal and IFN-stimulated conditions, we plotted the difference in transcript density ratio against the change in transcript abundance for 60 high-confidence transcripts (mean count>2 per cell in the IFN-stimulated sample) identified in the IFN-stimulated condition versus non-IFN control condition (Figure S4B). Most transcripts with increased condensate partitioning upon IFN stimulation were ISG-encoded (e.g., OASL, CMPK2, RSAD2, IRF7), while non-ISG transcripts (e.g., CLIP1, AKT3, IARS, ODC1) that were enriched in viral condensates at steady state showed reduced partitioning, suggesting an active exchange of cellular transcripts in viral condensates driven by ISG induction (Figure 4C).

These IFN-driven changes in RNA partitioning mirror the shift in RNA-binding dynamics inferred from the poly(A)-FISH experiments. We therefore proposed that RNA features favoring condensate partitioning at steady state are altered upon IFN stimulation. To test this hypothesis, we performed an RNA feature correlation analysis on 45 high-confidence, condensate-enriched transcripts (mean count per cell > 3, condensate density ratio > 1). Among eight RNA features analyzed (abundance; length; GC content; G-quadruplex count; and RNA folding of full transcript, 5’UTR, CDS, and 3’UTR), only the folding energy of the RNA 3’UTR showed a weak correlation with condensate partitioning under IFN stimulation (Spearman’s r = -0.3527, P = 0.02) (Figure 4D, Figure S4C). This contrasts with steady state, in which RNA folding of the CDS promotes partitioning (Figure 2E). Under IFN stimulation, more structured 3’UTRs favor condensate enrichment, likely mediated by IFN-inducible cellular RBPs in viral condensates, which interact with structured 3’UTRs in ISG transcripts.

We proposed these differences in RNA partitioning could, in part, explain why bats do not develop severe disease following Ebola virus infection compared to humans and other susceptible mammals. We therefore asked whether IFN-driven mRNA partitioning of bat transcripts differs from human transcripts. However, due to species-specific sequence variations, the human MERFISH probe panel could not be applied directly to fruit bat cells. Instead, we employed a computational approach to predict RNA folding free energies for 45 condensate-enriched transcripts encoded by bat genes. We then inferred their partitioning behaviors assuming that RNA feature preferences identified in human cells are conserved in bats. As controls, we performed the same RNA folding analysis on orthologs encoded by ferret and macaque, species known to develop severe disease after Ebola virus infection, like humans^62,63^. Because IFN responses typically induce fever in mammals, we included RNA folding energies predicted at 39°C to capture physiologically relevant changes in the RNA folding landscape. We observed overall lower RNA folding in bat, ferret, and macaque orthologs compared to human mRNAs at both 37°C and 39°C (linear regression slope <1) (Figure S5).

Although we did not observe a bat-specific bias in reduced RNA folding for the 45 high-confidence transcripts at 37°C, when we stratified these transcripts into IFN-induced (ISGs) and non-ISG categories based on experimentally measured changes in mRNA mean-count per cell with or without IFN induction, we found a notable difference: among mammalian ISGs, only bats exhibit overall reduced 3’UTR folding compared to their human orthologs. Ferret and macaque ISG transcripts have 3’UTR folding similar to their human counterparts. In contrast, non-ISG transcripts show no appreciable species-specific differences in 3’UTR folding across species (Figure 4E). Based on these findings, we predict ISG transcripts, which are enriched in viral condensates of IFN-stimulated human cells, are less likely to partition into viral condensates in bat cells.

Next, we tested whether 3’UTR RNA folding is sufficient to drive ISG mRNA partitioning in viral condensates using synthetic mScarlet reporters. To mimic endogenous ISG induction, we designed plasmids in which reporter transcription is driven by the Interferon Stimulation Response Element (ISRE) from human ISG15, enabling induction upon type 1 IFN stimulation. Upstream of the mScarlet CDS, we inserted the 5’UTR of human OASL, an ISG highly enriched in viral condensates that may support translation under IFN-stimulated conditions. Downstream, we tested three human ISG 3’UTRs (SAMHD1, IFIT1, and IFI6) with progressively lower predicted RNA folding and condensate partitioning measured by MERFISH. If 3’UTR folding is a primary determinant, the most structured 3’UTR (SAMHD1) should confer the highest condensate enrichment.

To test whether reduced 3’UTR folding enables transcripts to escape viral condensates in bat ISGs, we generated parallel reporters carrying Egyptian fruit bat orthologous 3’UTR. These bat 3’UTRs are predicted to be less structured than their human counterparts. All six reporters were detected using a common mScarlet RNA-FISH probe set (Figure 4F).

Upon IFN stimulation, reporters carrying bat, but not human 3’UTRs, follow the predicted relationship between RNA folding and condensate partitioning: the more folded the 3’UTR, the higher viral condensate partitioning (Figure 4F). Notably, the bat 3’UTRs were uniformly shorter than their human orthologs, suggesting that the relationship between 3’UTR folding and condensate partitioning may depend on RNA length. To test this hypothesis, we divided the 45 high-confidence transcripts enriched in viral condensates under IFN stimulation into short and long groups using 1,000 nt (median length) as a cutoff. Strikingly, the correlation between 3’UTR folding and condensate partition strengthened among shorter 3’UTRs but was lost among longer 3’UTRs (Figure 4G). This length dependence explains the reporter behavior: human IFI6, bat IFIT1, and bat IFI6 3’UTRs all fall below the 1,000 nt threshold and partition according to their predicted folding (Figure 4F). Within this regime, the less folded bat IFI6 3’UTR conferred lower condensate partitioning than its human counterpart.

Together, these findings reveal that upon type 1 IFN stimulation, Ebola viral condensates selectively and perhaps aberrantly enrich human ISG transcripts through 3’UTR folding dynamics, a process that may be attenuated in bats (Figure 4H).

### Cellular mRNA partitioning in viral condensate during authentic Ebola infection

To confirm cellular mRNA partitioning in the presence of all viral proteins, we quantified IFIT3 (ISG) transcript localization in viral condensates during authentic Ebola virus infection using a biologically contained Ebola virus, constructed by replacing the critical viral transcription factor VP30 with GFP, restricting replication to VP30-expressing cells^64^. Following type I IFN stimulation at 6 h post infection, IFIT3 mRNA was detected in viral condensates across multiple infection states over a 24 h time course, including early infection (GFP−; small condensates), active transcription (GFP+), replication (vRNA+ accumulation), and late infection with peripheral nucleocapsid filaments (Figure 5A).

**Figure 5.**
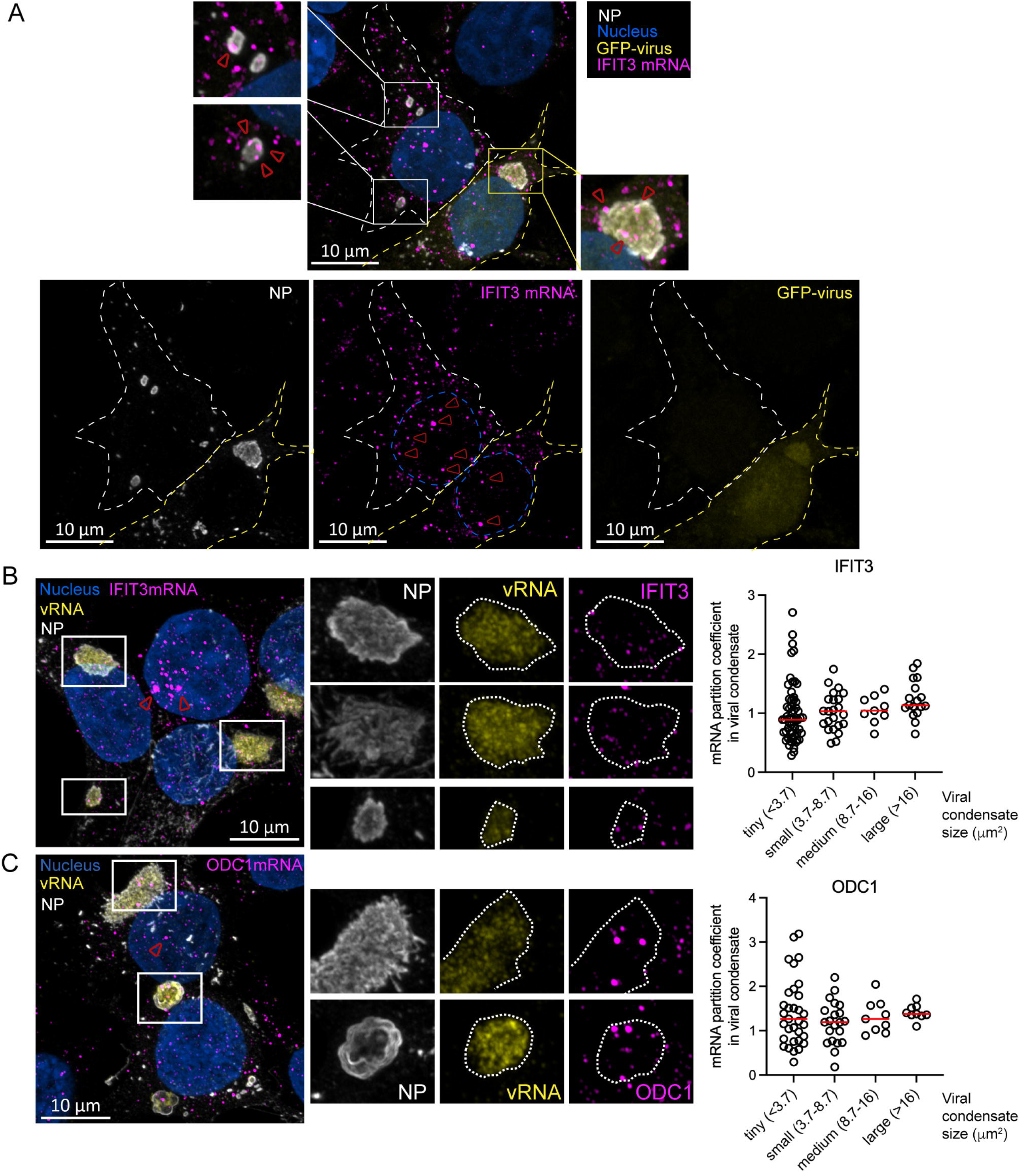
Cellular mRNA partitioning in viral condensate during Ebola virus infection. **A.** Confocal microscopy of fixed and stained HEK293T cells transfected with Ebola VP30 and subsequently infected with Ebola-dVP30-GFP virus at MOI=5. Infected cells were incubated with type 1 IFN (1000U/ml) at 6-hour post infection and fixed at 24-hour post infection. Viral condensates are marked by NP immunofluorescence staining. Endogenous IFIT3 mRNA labeled by SABER-FISH. Nuclei stained with Hoechst. Virus infected cells marked by GFP reporter. Scale bar: 10 µm. **B. C.** Quantification of endogenous IFIT3 mRNA (**B**) and ODC1 mRNA (**C**) partition coefficient in replication-active viral condensates in Ebola-infected cells measured by SABER-FISH. Replication-active viral condensates detected by enriched viral genome (vRNA). Median values with individual data points (per condensate) from two experiments are shown (with 32 cells for IFIT3 FISH and 27 cells for ODC1 FISH analyzed).

Compared with NP+VP35 reconstitution and minigenome systems, infection revealed additional regulatory context. IFN-induced IFIT3 transcription was markedly reduced in infected cells, consistent with VP24-mediated inhibition of STAT1 nuclear translocation and downstream ISG activation^65,66^, as evidenced by diminished nuclear transcriptional bursts relative to neighboring uninfected cells (Figure 5B). Despite this suppression and cell-to-cell heterogeneity in IFN responses, some IFIT3 transcripts produced during infection still partitioned into viral condensates, indicating that condensate-mediated ISG recruitment persists under viral immune antagonism.

Notably, IFIT3 enrichment increases with condensate size (Figure 5B). A similar trend was observed for non-ISG ODC1 (Figure 5C), suggesting host mRNA recruitment may contribute to condensate growth, potentially stabilizing replication compartments during infection. Together, these results validate mRNA partitioning observed in reconstituted systems also occurs during infection and may support viral replication compartment stability.

### Functional Consequences of Cellular mRNA Partitioning into Viral Condensates

We next examined the functional impact of cellular mRNA partitioning into Ebola viral condensates; specifically, we asked how sequestration into condensates might influence mRNA stability and protein synthesis. Given that viral condensates can spatially exclude cellular machinery such as ribosomes^67,68^, we determined how mRNA partitioning correlates with transcript and protein accumulation.

We quantified mRNA and protein levels for representative cellular transcripts with varying degrees of condensate enrichment, as determined by MERFISH. For mRNA quantification, we selected five transcripts spanning a range of partition coefficients in steady-state and IFN-stimulated HEK293T cells. For protein analysis, we sampled three transcripts per condition, limited to proteins with commercially available antibodies suitable for western blotting. We compared mRNA and protein levels between non-transfected cells and cells co-expressing increasing levels of viral condensate scaffold proteins NP and VP35 to correlate changes in cellular transcript and protein accumulation with viral condensate formation.

At steady state (without IFN stimulation), transcripts that are enriched in viral condensates (e.g., CCDC92, ODC1, EIF2AK2) display a modest increase in mRNA abundance (up to 1.5-fold), proportional to the level of viral NP and VP35 expressed to reconstitute condensates. In contrast, transcripts not enriched in condensates (SAT1, MOV10) show no significant changes in mRNA level with or without of NP and VP35 (Figure 6A). We observe a different effect on the proteins encoded by these transcripts. For transcripts enriched in viral condensates, although mRNA increased, protein expression from those transcripts (DCP1A, ODC1, EIF2AK2) decreased, suggesting that sequestration into condensates impedes their translation (Figure 6B).

**Figure 6.**
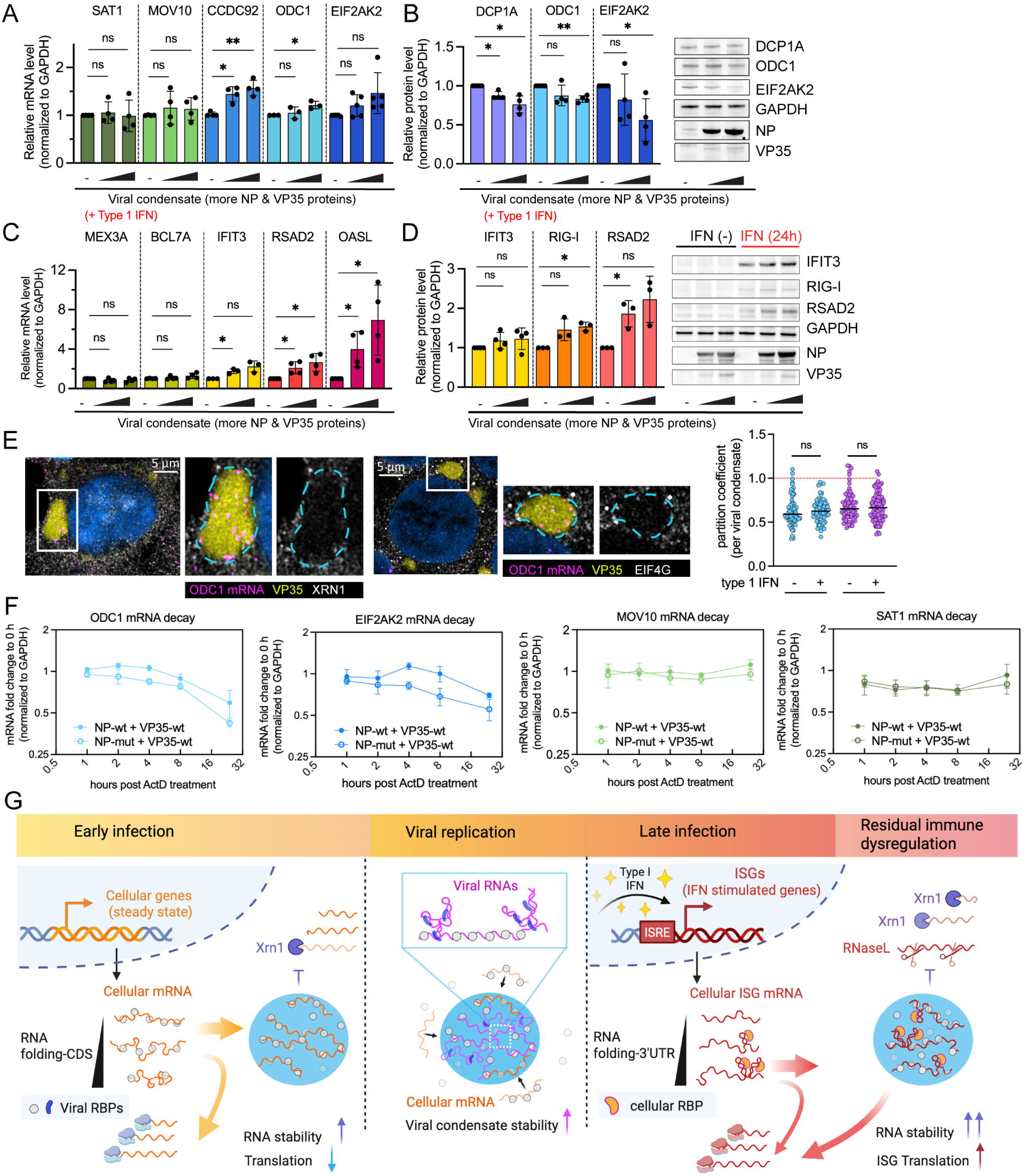
Partitioning in Ebola virus condensates changes RNA stability and protein accumulation. **A.** Relative abundance of five cellular mRNAs, ordered from least to most enriched in the viral condensate at steady state. **B.** Relative abundance of three cellular proteins, ordered from the least to most enriched corresponding transcript in the viral condensate at steady state. **C.** Relative abundance of five cellular mRNAs, ordered from least to most enriched in the viral condensate upon IFN stimulation. **D.** Relative abundance of three cellular proteins, ordered by corresponding transcripts enrichment in the viral condensate upon IFN stimulation. Oligo-dT-primed RT-qPCR and western blot analyses on total RNA/protein extracted from HEK293T cells with increasing level of NP- and VP35-driven condensate formation. GAPDH mRNA and protein level were used for normalization. Representative western blots for each protein target, loading control, and viral condensate scaffold NP and VP35 are shown. Unstimulated control western blots (IFN-) indicate basal level expression of ISG proteins and are not used for quantification. Mean±SD of individual data points from four experiments are compared. **: P<0.01, *: P<0.05, ns: not significant, by paired t test. **E.** Confocal microscopy of fixed and stained HEK293T (human kidney) cells co-expressing untagged Ebola virus NP and mNG-tagged Ebola virus VP35. Boxed condensate within a merge overview is magnified to show individual channels for VP35-labeled viral condensates, XRN1/EIF4G staining, and ODC1 mRNA. Nuclei stained with Hoechst and shown in blue. Scale bar: 5 µm. Endogenous ODC1 mRNA was labeled by SABER-FISH, while endogenous XRN1 or EIF4G were visualized by immunofluorescence. Partition coefficients of XRN1 or EIF4G in ODC1 mRNA-containing viral condensates are shown. Median and individual data points from two experiments (>10 cells in each condition per experiment) are plotted and compared. ns: not significant by Kolmogorov-Smirnov test. **F.** RNA half-life analysis was performed to monitor representative condensate-enriched cellular mRNAs (ODC1 and EIF2AK2) and not enriched (MOV10 and SAT1) in the absence of IFN simulation. Actinomycin D (ActD) was added to cells at 0 hour (10 ug/ml) to block transcription, one day after transfecting cells to reconstitute viral condensate with HA-mNG-tagged Ebola VP35 (VP35-wt) plus either wild type (NP-wt) or RNA-binding mutant of NP (NP-mut). Relative mRNA levels of each target were quantified by rt-qPCR and normalized to the reference gene (GAPDH). For each target in each condition, the mRNA level at all time points (1, 2, 4, 8, 24 hours) post-ActD addition was normalized to the starting level measured before ActD addition (0 hour) and plotted. Results of four experiments, with the mean value of four technical replicates for each experiment are plotted as mean±SEM. **G.** A proposed model describing context-dependent principles for viral condensates recruiting cellular and viral RNAs and the downstream functional impact on RNA stability and translation. Created using BioRender.

Upon type I IFN stimulation, transcripts enriched in viral condensates (e.g., IFIT3, RSAD2, OASL) display more robust increases in mRNA accumulation, with OASL showing a >6-fold increase. Control transcripts not enriched in condensates (MEX3A, BCL7A) remain unchanged in mRNA accumulation (Figure 6C). IFN-stimulated cellular proteins (RIG-I and RSAD2) increase in proportion to the amount of viral condensate scaffold proteins expressed, suggesting that condensate partitioning under IFN-stimulated conditions enhances both mRNA and protein accumulation of cellular ISGs (Figure 6D).

Viral condensates may promote mRNA stabilization by excluding canonical RNA decay machinery such as the 5’-3’ exonuclease XRN1^69^. Indeed, XRN1 is partially excluded from viral condensates, with a median partition coefficient of 0.6 under both steady state and IFN-stimulated conditions (Figure 6E). To validate this mechanism, we analyzed mRNA degradation kinetics after Actinomycin D treatment (transcription inhibition) for condensate-enriched transcripts (ODC1, EIF2AK2) under wild-type and RNA-binding mutant NP conditions. In condensates formed by wild-type RNA-binding NP, both condensate-enriched transcripts consistently retain higher abundance across multiple timepoints compared with condensates formed by NP unable to bind RNA. Control transcripts (SAT1, MOV10), which are not enriched in viral condensates, retain equivalent levels in wild-type and RNA-binding mutant condensates across multiple timepoints (Figure 6F). These results suggest that partitioning into viral condensates protects mRNAs from XRN1-mediated degradation.

In contrast to this +/- IFN-independent mRNA stabilization, we examined the IFN-dependent effect of condensate partitioning on protein translation. Previous studies have shown that Ebola viral condensates exclude ribosomes during early stages of infection^68,70^. Consistent with this observation, we found EIF4G, a key translation initiation factor^71^, displayed reduced partitioning (median coefficient = 0.65) into viral condensates, supporting the notion that condensate-resident mRNAs are less accessible to canonical translation machinery. This sequestration likely contributes to the modest reduction in protein accumulation observed for condensate-enriched transcripts at steady state. However, EIF4G remains partially excluded from viral condensates under IFN-stimulated conditions, suggesting viral condensates remain translationally repressed (Figure 6E).

During IFN responses, while global translation is attenuated, ISG mRNAs are selectively translated via alternative pathways^72^. We propose this selective translational mechanism enables ISG mRNAs to overcome condensate-imposed translational repression. An alternative, non-mutually exclusive explanation is that Ebola VP35-mediated inhibition of the IFN-induced translation inhibitor PKR contributes to de-repression of ISG translation^73,75^. In this context, the combined effects of enhanced mRNA stability within viral condensates and preferential translation of ISG mRNAs, likely occurring downstream of condensate partitioning, result in a net increase in ISG protein levels upon IFN stimulation (Figure 6D).

## Discussion

RNA viruses that impact over a billion people annually—including measles virus, influenza A virus, and RSV—form phase-separated, intracellular condensates during infection^4–6,10^. While these viral condensates are recognized as a mechanism to concentrate viral building blocks for replication, their potential to interact with host cellular RNAs and influence host gene expression remain elusive. Our work quantitatively mapped the partitioning of over 100 disease-relevant cellular mRNAs in Ebola viral condensates. We then illuminated the molecular principles governing RNA partitioning into viral condensates, dissected their functional impact on virus and host evolution, and revealed hidden links between condensate function and viral disease pathogenesis.

We discovered Ebola virus–induced condensates can selectively partition cellular mRNAs based on distinct molecular features and modulate RNA fate in a IFN-dependent manner. Using poly(A)-FISH combined with RNA-binding mutants of viral proteins, we delineated the molecular basis of RNA recruitment by condensates in human and fruit bat cells. Given the conserved, sequence-independent, RNA-binding properties of nucleoproteins across negative-strand RNA viruses^74,76,77^, we propose that host RNA partitioning into viral condensates may represent a common feature for this group of viruses. Indeed, a recent study using *in situ* sequencing with photoisolation chemistry (PIC) detected cellular RNAs within viral condensates in mumps virus– infected cells^78^, supporting the generality of this phenomenon. However, limitations in spatial resolution and quantification in the PIC-based approach—stemming from photoirradiating only regions of interest for downstream sequencing—preclude robust analysis of the relative enrichment of cellular transcripts in viral condensates at scale^79^, which is necessary to discern RNA features mediating subcellular localization. In contrast, MERFISH-based subcellular transcriptomics enables precise, single-cell quantification of condensate-associated transcripts *in situ*. This technique enabled us to reveal viral condensates preferentially recruit RNAs with minimally folded coding regions at steady state and RNAs with highly structured 3’UTRs following type I interferon (IFN) stimulation. We further validated the correlation between RNA structural features and condensate-partitioning behavior using rationally designed synthetic reporter RNAs and antisense oligo blockers targeting endogenous mRNA in cells. This context-dependent partitioning echoes observations in endogenous cellular condensates, such as P-bodies^80^, and aligns with a recent transcriptome-wide study showing RNA structure influences localization to endogenous condensates^81^. Together, our findings uncover a previously unrecognized layer of virus–host interaction, in which viral condensates selectively partition host mRNAs in a context- and RNA folding-dependent manner.

Importantly, our findings suggest viral condensates can stabilize resident RNAs, a clear advantage for viruses. As specialized compartments for viral RNA synthesis, condensates that prolong viral RNA persistence without requiring enhanced polymerase activity may confer an evolutionary benefit. Consistent with this idea, previous studies have shown rabies virus condensates contain RNAs that are resistant to exogenous RNase A treatment in fixed cells^82^, suggesting physical protection. Condensate-mediated RNA stabilization could offer a compelling molecular basis for viral persistence, a phenomenon well documented in Ebola survivors^83,84,86^. However, this stabilizing effect is not exclusive to viral RNA. Host mRNAs with features favoring condensate partitioning, such as reduced coding region folding or highly structured 3’UTR, are also preserved beyond their intrinsic lifetime. This has consequences for ISGs, which require tight calibration to prevent excessive immune activation^85,87^. We found a subset of ISG transcripts involved in innate immune response and regulation are aberrantly stabilized and translationally upregulated within Ebola viral condensates. Such prolonged ISG expression may contribute to persistent inflammation, a phenomenon well documented in EVD survivors^24,88,89^. By linking viral condensate-mediated RNA stabilization to immune dysregulation, our study offers a new molecular framework for understanding the chronic immunopathology that can follow acute RNA virus infections.

The molecular features that govern RNA partitioning into viral condensates may have broad evolutionary implications for viruses and hosts. We found several RNA viruses including measles virus, rabies virus and RSV, previously known to form condensates^46,90,91^, share a common preference for reduced secondary structure within their protein-coding sequences. This suggests evolutionary pressures constrain viral sequence variation to preserve a minimally structured genome architecture compatible with condensate residency. On the host side, we identified a subset of ISGs with divergent 3’UTR folding across mammalian species with distinct outcomes following Ebola virus infection. Notably, in fruit bats, which are reservoir hosts for multiple high-consequence RNA viruses, the 3’UTRs of cellular ISG transcripts are less structured than their human counterparts. We propose this reduced 3’UTR folding liberates bat ISG transcripts from sequestration into viral condensates, thereby allowing normal function and turnover of these antiviral transcripts in bat cells. Such a mechanism may contribute to the uniquely balanced and nonpathogenic antiviral responses observed in bats^27,92^. Through comparative RNA structure analysis, our findings suggest an evolutionary arms race over RNA structure and its recruitment to viral condensates.

Based on our findings with reconstituted viral condensates and validation with authentic infection, we propose Ebola viral condensates can dynamically partition viral and cellular RNA throughout infection (Figure 6G). Early in infection, condensate formation is driven by an accumulation of viral proteins and stabilized in part by host mRNAs with minimally folded coding regions. These host transcripts may help stabilize the viral condensate precursor, which is essential for the virus to establish productive infection in the host^93^. As infection progresses beyond Ebola virus protein-mediated immune evasion^94^, type I IFN response triggers ISG expression, which unleashes IFN-stimulated ribonucleases such as RNase L to degrade viral and host mRNA and leads to condensate partitioning of IFN-stimulated dsRNA-binding proteins such as PKR and ADAR^58,60^. This shift promotes selective recruitment of ISG transcripts with structured 3’UTRs, enhancing their transcripts and protein accumulation. While future studies are needed to determine whether persistent and aberrant ISG expression driven by viral condensate partitioning contributes directly to immunopathology following acute infection in physiologically relevant systems, our findings underscore the importance of developing therapeutic strategies to disrupt and eliminate viral condensates, not only to limit viral replication, but also to mitigate long-term immune dysregulation associated with infection.

### Limitations of the study

While cellular mRNA partitioning was measured at a single post-transfection time point (24 hours) to allow formation of viral condensates, and using a selected panel of 136 cellular genes, chosen for their relevance to Ebola infection and innate immunity; genome-wide and time-resolved studies may uncover additional principles and dynamics governing cellular transcriptome remodeling. Due to the usage of HEK293T, a cell line that has low sensitivity to IFN stimulation, the dosage and timing used in our experiments to efficiently induce ISG transcription could be higher and longer than the typical range needed for other cells and tissues. We acknowledge that our findings in Egyptian fruit bat (*Rousettus aegyptiacus*) should not be generalized to bats broadly. We selected *R. aegyptiacus* as a useful comparative model for bat-filovirus interactions because cell lines from this species are among the few established for filovirus research^97–100^, are amenable to transient transfection and permissive to Ebola virus infection *in vitro*^99,101^, and because the Rousettus bat is a known reservoir of Marburg virus^40^. All RNA folding analyses performed in this study are sequence-based computational predictions, which do not account for the contribution of RNA-binding proteins and chemical modifications to RNA structures in living cells. We focused on thermodynamic ensemble free energy because it provides a thermodynamic quantity for considering RNA folding and unfolding under different physiological conditions, such as elevated body temperature. Other relevant parameter describing RNA folding, including ensemble diversity, should be examined in future studies. Finally, while minimal coding-region folding appears conserved in the viral genome across negative-strand RNA viruses, RNA features selected for condensate partitioning may differ across other RNA viruses, host cell types, and host species.

## Resource Availability

### Lead Contact

Requests for further information should be directed to the lead contact, Erica Ollmann Saphire.

### Materials availability

All materials and resource request should be directed to and will be fulfilled by the lead contact.

### Data and code availability

Data in this manuscript will be shared by the lead contact upon requests. The scripts used for the computational analyses in the manuscript, including MERFISH-based subcellular spatial transcriptomics analysis of host mRNA partitioning into Ebola virus condensates, RNAfold-based RNA secondary structure analysis, and additional transcript feature analyses were deposited at Zenodo (https://doi.org/10.5281/zenodo.23019809).

## Acknowledgments

We thank Ayana Adams (a previous member of the Saphire Lab) for technical assistance in molecular cloning and plasmid preparation; Sarah Van Tol and Vincent Munster (NIH) for sharing immortalized bat kidney cells RAKSM line; Lennard W. Wiesner for sharing an optimized SABER-FISH protocol; Alexander Ploss for sharing Vero-E6 cells, and Princeton Research Computing resources at Princeton University. We thank Yoshihiro Kawaoka and Peter Halfmann (University of Wisconsin) for sharing the EBOV-GFP-ΔVP30 system with the Saphire lab. We thank Amy R. Strom and Lindsay A. Becker (Brangwynne lab), Karishma Bisht (te Velthuis lab), and Sarah Van Tol (NIH) for helpful feedback and discussion of the manuscript/project. MERFISH experiments were supported by the 2024 Tullie and Rickey Families SPARK Award for Innovations in Immunology to J.F. [La Jolla Institute for Immunology (LJI)]. This work was supported by the Howard Hughes Medical Institute, the Princeton Biomolecular Condensate Program, the Princeton Center for Complex Materials, a MRSEC (NSF DMR-2011750), the St Jude Collaborative on Membraneless Organelles, the AFOSR MURI (FA9550-20-1-0241) and the Chan Zuckerberg Initiative Exploratory Cell Network (to C.P.B.), a National Institutes of Health grant DP2 AI175474-01 (to A.J.W.t.V.). G.W.Y. acknowledges funding by the UC San Diego Sanford Stem Cell Institute. EOS is an investigator at LJI, which receives annual institutional support from Kyowa Kirin, Inc. (KKUS). The funding received from KKUS does not specifically support the research described in this publication. J.F. is supported by the 2025 Omenn Darling Postdoctoral Fellowship through Princeton University.

## Author contributions

J.F. conceptualized this project, including methodology, investigation, formal analysis, and interpretation, with supervision from E.O.S., J.F. performed experiments and analysis, D.L. and C.K.M. were supervised by G.W.Y., developed the computational pipeline and performed subcellular MERFISH data processing, performed RNA features prediction, and drafted methods in the corresponding section. R.W. performed biologically contained Ebola virus (EBOV-GFP-ΔVP30) infection in BSL2+ containment. H.M.T. with advice from C.P.B., performed live cell imaging and image analysis for hypotonic dissolution assay, drafted methods and plotted data in the corresponding section. T.C. assisted with cellprofiler-based image analysis pipeline. Y.C. and C.C. performed all MERFISH imaging, generated spatial data of detected transcripts, and drafted the corresponding methods. J.F. wrote the original draft and generated all figures, with substantial edits from E.O.S. and A.J.WtV. J.F., E.O.S., A.J. WtV, C.P.B.,

G.W.Y. acquired funding. All authors reviewed and edited the manuscript.

## Declaration of generative AI and AI-assisted technologies in the writing process

During the preparation of this work the author used ChatGPT to improve the clarity and flow of writing. After using this tool, the author reviewed and edited the content as needed and takes full responsibility for the content of the publication.

## STAR METHODS

### Key resources table

| REAGENT or RESOURCE | SOURCE | IDENTIFIER |
| --- | --- | --- |
| Antibodies |  |  |
| Rabbit, anti-EBOV NP | GeneTex | GTX134031 |
| Mouse, anti-EBOV VP35<br>Clone 6C5 | Kerafast | KF-Ab01229-2.0 |
| Human, anti-Ebola NP<br>Clone KZ51 | Absolute antibody | Ab00692-10.0 |
| Vizgen Protein Stain Anti-mouse | Vizgen | 20300101 |
| Vizgen Protein Stain Anti-human | Vizgen | 20300105 |
| anti-EIF4G mouse monoclonal antibody (A-10) | Santa Cruz Biotechnology | sc-133155 |
| anti-XRN1 mouse monoclonal antibody (C-1) | Santa Cruz Biotechnology | sc-165985 |
| anti-GAPDH mouse monoclonal antibody (6C5) | Santa Cruz Biotechnology | sc-32233 |
| anti-DCP1A rabbit monoclonal antibody | Abcam | ab183709 |
| anti-ODC1 rabbit polyclonal antibody | Proteintech | 28728-1-AP |
| anti-IFIT3 rabbit polyclonal antibody | Sigma-Aldrich | HPA059914 |
| anti-HA (C29F4) rabbit monoclonal antibody | Cell Signaling Technology | 3724T |
| anti-HA Affinity Matrix (from rat IgG1) | Roche | 11815016001 |
| anti-EIF2AK2/PKR rabbit monoclonal antibody | Abcam | ab184257 |
| anti-RIGI mouse monoclonal antibody Clone 4G1B6 | Thermo Fisher Scientific | MA5-31715 |
| anti-RSAD2 rabbit polyclonal antibody | Proteintech | 28089-1-AP |
| Goat anti-Rabbit Alexa Fluor 647 | ThermoFisher Scientific | A-21245 |
| Peroxidase AffiniPure Goat anti-Rabbit IgG (H+L) | Jackson ImmunoResearch | 111-035-144 |
| Peroxidase AffiniPure Goat anti-mouse IgG (H+L) | Jackson ImmunoResearch | 115-035-062 |
| Bacterial and virus strains |  |  |
| EBOV-ΔVP30-GFP | (Halfmann et al. 2008)<sup>64</sup> | N/A |
| Chemicals, peptides, and recombinant proteins |  |  |
| DMEM high glucose, pyruvate | Thermo Fisher Scientific | 11995065 |
| DPBS, no calcium, no magnesium | Thermo Fisher Scientific | 14190144 |
| Opti-MEM reduced serum medium | Thermo Fisher Scientific | 31985062 |
| Penicillin-Streptomycin | Thermo Fisher Scientific | 15140122 |
| Hoechst 33342 | Thermo Fisher Scientific | 62249 |
| Deionized Formamide | Thermo Fisher Scientific | AM9342 |
| ProLong Gold Antifade Mountant | Thermo Fisher Scientific | P36934 |
| Lipofectamine 3000 | Thermo Fisher Scientific | L3000015 |
| Halt™ Protease inhibitor cocktail and phosphatase inhibitor cocktail | Thermo Fisher Scientific | 78445 |
| Invitrogen Ultrapure BSA | Thermo Fisher Scientific | AM2616 |
| NuPage™ LDS sample buffer (4X) | Thermo Fisher Scientific | NP0007 |
| Invitrogen TRIzol Reagent | Thermo Fisher Scientific | 15596026 |
| Invitrogen TRIzol LS Reagent | Thermo Fisher Scientific | 10296010 |
| Paraformaldehyde (PFA) | Electron Microscopy Sciences | 15710 |
| Fetal Bovine Serum | Atlanta biological | S11150H |
| Human Fibronectin Plasma | Sigma-Aldrich | F0895 |
| Recombinant Universal Type 1 IFN | PBL Assay Science | 11200 |
| Ammonium persulfate | Sigma-Aldrich | 09913-100G |
| TEMED | Sigma-Aldrich | T7024-25 ML |
| Actinomycin D | Sigma-Aldrich | A4262 |
| Benzonase nuclease | Sigma-Aldrich | E8263 |
| SSC buffer | Sigma-Aldrich | S6639 |
| Dextran Sulfate, sodium salt | Millipore Sigma | 265152 |
| Tween-20 | Millipore Sigma | P7949 |
| Proteinase K | NEB | P8107S |
| RNase inhibitor | NEB | M0314L |
| Clarity Western ECL substrate | Bio-Rad | 1705061 |
| TransIT-LT1 transfection reagent | Mirus Bio | MIR2300 |
| Stellaris® RNA FISH Hybridization Buffer | LGC, Biosearch Technologies | SMF-HB1-10 |
| Stellaris® RNA FISH Wash Buffer A | LGC, Biosearch Technologies | SMF-WA1-60 |
| Stellaris® RNA FISH Wash Buffer B | LGC, Biosearch Technologies | SMF-WB1-20 |
| Blocking Buffer C Premix | Vizgen | 20300100 |
| Formamide Wash Buffer | Vizgen | 20300002 |
| Gel Embedding Premix | Vizgen | 20300004 |
| MERSCOPE Sample prep kit | Vizgen | 10400012 |
| Clearing Premix | Vizgen | 20300003 |
| Sample Prep Wash Buffer | Vizgen | 20300001 |
| Poly T Reagent | Vizgen | 20300021 |
| Applied Biosystems™ PowerUp™ SYBR™ green master mix | Thermo Fisher Scientific | A25742 |
| NuPAGE™ Bis-Tris Mini Protein Gels, 4-12%, (1.5 mm, 10-well) | Thermo Fisher Scientific | NP0335BOX |
| Critical commercial assays |  |  |
| Direct-zol RNA miniprep kit | ZymoResearch | R2050 |
| High-capacity cDNA Reverse Transcription kit | Thermo Fisher Scientific | 4368814 |
| NEBuilder® HiFi DNA Assembly Master Mix | NEB | E2621X |
| Deposited data |  |  |
| Human reference genome | Ensembl | GRCh38.p14 |
| Ferret reference genome | NCBI | ASM1176430v1.1<br>(GCF_011764305.1) |
| Macaque reference genome | NCBI | Mmul_10<br>(GCF_003339765.1) |
| Egyptian fruit bat reference genome | NCBI | mRouAeg1.p<br>(GCF_014176215.1) |
| Experimental models: Cell lines |  |  |
| HEK293T | ATCC | RRID: CVCL_0063 |
| Vero E6 | ATCC, a kind gift from Alexander Ploss | RRID: CVCL_0574 |
| Huh7 | A kind gift from Mart M. Lamers | RRID: CVCL_0336 |
| RaKSM-2.5i | A kind gift from Vincent Munster | RRIF: CVCL_YZ04 |
| Oligonucleotides |  |  |
| T30-Quasar 570 FISH probe | LGC, Biosearch Technologies | T30-Quasar 570-1 |
| Oligo(dT)<sub>20</sub> primer | ThermoFisher | 18418020 |
| SABER-FISH probes, see Table S2 | This paper | N/A |
| qPCR primers, see Table S3 | PrimerBank | N/A |
| LNA/DNA mixer ASO targeting human ODC1:<br>+C*+T*CTG+TG+GCGT+T<br>TCA*+T*+C (*:PS-bond;<br>+:LNA) | This paper | N/A |
| Recombinant DNA |  |  |
| pCAGGS-NP | (Fang et al. 2023; Schmidt et al. 2018)<sup>35,56</sup> | N/A |
| pCAGGS-NPssRNAmut | This paper | N/A |
| pCAGGS-HA-mNG-VP35 | (Fang et al. 2023)<sup>35</sup> | N/A |
| pCAGGS-HA-mNG-VP35-dsRNAmut | This paper | N/A |
| pCAGGS-VP30 | (Fang et al. 2023; ;<br>Schmidt et al. 2018)<sup>35,56</sup> | N/A |
| pCAGGS-T7pol | (Fang et al. 2023; Schmidt<br>et al. 2018)<sup>35,56</sup> | N/A |
| pCAGGS-L | (Fang et al. 2023; Schmidt<br>et al. 2018)<sup>35,56</sup> | N/A |
| pT7-2cis-MG | (Fang et al. 2023)<sup>35</sup> | N/A |
| pCAGGS-huODC1_5'UTR-mScar_Ori_huODC1_3'UT<br>R | This paper | N/A |
| pCAGGS-huODC1_5'UTR-mScar_Mod1_huODC1_3'<br>UTR | This paper | N/A |
| pCAGGS-huODC1_5'UTR-mScar_Mod2_huODC1_3'<br>UTR | This paper | N/A |
| pC3_ISG15-ISRE_huOASL5'UTR_mS<br>carMod1_huSAMHD1_3'U<br>TR | This paper | N/A |
| pC3_ISG15-ISRE_huOASL5'UTR_mS<br>carMod1_huIFIT1_3'UTR | This paper | N/A |
| pC3_ISG15-ISRE_huOASL5'UTR_mS<br>carMod1_huIFI6_3'UTR | This paper | N/A |
| pC3_ISG15-ISRE_huOASL5'UTR_mS<br>carMod1_batSAMHD1_3'<br>UTR | This paper | N/A |
| pC3_ISG15-ISRE_huOASL5'UTR_mS<br>carMod1_batIFIT1_3'UTR | This paper | N/A |
| pC3_ISG15-ISRE_huOASL5'UTR_mScarMod1_batIFI6_3'UTR | This paper | N/A |
| Software and algorithms |  |  |
| CellProfiler v4.2.6 | Broad Institute | RRID: SCR_007358 |
| Cellpose v2.3.2 | (Stringer et al. 2021)<sup>95</sup> | RRID: SCR_021658 |
| Fiji(ImageK2) | NIH | RRID: SCR_003070 |
| Python v3.10 | Python Software Foundation | RRID: SCR_008394 |
| MERlin | Vizgen | N/A |
| SpatialDat-io | scverse | N/A |
| Sopa | scverse | N/A |
| Bento | Bento-tools<sup>96</sup> | N/A |
| OpenCV | OpenCV | RRID:SCR_015526 |
| scikit-image | scikit-image | RRID:SCR_021142 |
| Shapely | Shapely | N/A |
| GeoPandas | GeoPandas | N/A |
| VinneRNA (RNAfold) | VinneRNA (RNAfold) | RRID:SCR_008550 |
| QGRS Mapper | (Kikin et al. 2006)<sup>102</sup> | N/A |
| GraphPad Prism 10 | GraphPad Software | RRID:SCR_002798 |
| Nikon NIS-Elements | Nikon | N/A |
| Zen | Zeiss | N/A |
| **Other** |  |  |

## EXPERIMENTAL MODEL AND STUDY PARTICIPANT DETAILS

### Cell cultures

Human embryonic kidney cells HEK 293T, female origin (RRID:CVCL_0063), Human hepatoma cell line Huh7, male origin (RRID:CVCL_0336), immortalized Egyptian fruit bat kidney cells RaKSM-2.5i, or RAKSM, sex origin not reported (RRID:CVCL_YZ04), and immortalized African green monkey kidney cells VeroE6, female origin, were maintained in high glucose, pyruvate (Thermofisher, 11995065) supplemented with 10% fetal bovine serum (FBS) (Atlanta Biological, S11150H), 100 U/mL GIBCO^TM^ penicillin-streptomycin (Thermofisher, 15140122), and grown at 37°C and 5%CO_2_.

The biologically contained Ebola virus, EBOV-ΔVP30-GFP, was generated in HEK 293T cells^64^.

## METHOD DETAILS

### PolyA-FISH sample preparation, imaging, and image quantification

HEK293 T cells (3-4X10^4^ cells/well) were seeded in human fibronectin (Sigma, F0895) coated glass-bottom IBIDI µ-slide 8 well high glass bottom (IBIDI, 80807) 24 hours prior to transfection. RAKSM cells (2×10^4^ cells/well) were seeded directly in IBIDI slides without fibronectin. Cells were co-transfected with 50 ng/well of pCAGGS-NP or pCAGGS-NPssRNAmut paired with 50 ng/well pCAGGS-HA-mNG-VP35 or pCAGGS-HA-mNG-VP35-dsRNAmut using Trans IT LT-1 transfection reagent (Mirus Bio, MIR2300) for HEK293T cells or Lipofectamine^TM^ 3000 transfection reagent (Thermofisher, L3000015) for RAKSM cells. For IFN-stimulated samples, at 4 hours post-transfection, cells were incubated with complete cell culture media-containing 1000 U/ml recombinant universal type 1 IFN (PBL assay science,11200) for 24 hours. At 1 day post transfection or 24 hours post-IFN treatment, samples were chemically fixed with 4%PFA in PBS for 10-15 minutes and permeabilized with 70% ethanol at 4°C overnight or for several days to ensure complete permeabilization. “Stellaris™ RNA FISH Probes recognized and labelled with Quasar™ 570 dye (LGC, Biosearch Technologies, T30-Quasar 570-1) were hybridized to polyA containing RNAs, following the manufacturer’s instructions available online at www.biosearchtech.com/stellarisprotocols.

Briefly, samples were hybridized with 125 nM T30-Quasar 570 FISH probes and 1:1000 diluted anti-Ebola virus NP polyclonal antibody (GeneTex, GTX134031) in Stellaris® RNA FISH Hybridization Buffer (LGC, Biosearch Technologies, SMF-HB1-10) containing 10% deionized formamide (Thermofisher, AM9342) at 37°C for 4 hours. Secondary antibody incubation using Invitrogen Goat anti-Rabbit AlexaFluor^TM^647 (Thermofisher, A-21245) was combined into the second wash with Stellaris® RNA FISH Wash Buffer A (LGC, Biosearch Technologies, SMF-WA1-60) incubation at 37°C for 30 minutes. Nuclei were stained with Hoechst 33342 (Thermo Fisher, 62249). Samples were washed with Stellaris® RNA FISH Wash Buffer B and mounted in ProLong^TM^ Gold anti-fade mountant (Thermo Fisher, P36934) and the next day imaged on Zeiss LSM980-airyscan system with a 60x oil objective using airyscan-MULTI mode.

To quantify the size and number of viral condensates, an image analysis pipeline was built in CellProfiler 4.2.6^103^, which uses the mNG-VP35 channel for viral condensate segmentation, Hoechst channel for nuclei segmentation, and either unmodified polyT FISH channel or in the case of mutant with mRNA staining depleted from viral condensates, polyT-FISH combined with NP channel for cytoplasm segmentation. Cytoplasm segmentation for human cells was performed using Cellpose 2.3.2^95^ module; for bat cells, segmentation was instead performed using the identify secondary object module with nuclei as the primary object. Mean intensities of polyT-FISH signal in viral condensates and in cytoplasm excluding viral condensates were used to quantify the mRNA partition coefficient per viral condensate.

### Hypotonic dissolution and image quantification of fixed and stained cells

HEK293T cells were seeded, transfected and stained for viral protein NP, cell nuclei, and endogenous polyA-containing mRNAs as described in the previous section. Hypotonic shock was performed by incubating live cells in media with two times diluted in nuclease-free water for 15 minutes at 37°C incubator prior to fixation. To quantify coefficient of variation (CV) of mNG-VP35 marked viral condensates in different conditions, an image analysis pipeline was built in CellProfiler as described above, which uses Hoechst channel for nuclei segmentation and unmodified polyT FISH signal for cytoplasm segmentation. MeanIntensity and StdIntensity of the mNG-VP35 channel for each segmented cell were used for CV quantification.

### Hypotonic dissolution assay and image quantification in live cells

HEK293T cells were seeded, transfected as described in the previous section and stained for cell nuclei prior to live cell imaging. A Nikon spinning disk confocal microscope system, equipped with a Yokogawa W1 SoRa spinning disk confocal scanhead and built around a Nikon Ti2-E fully motorized microscope harboring dual Hamamatsu Fusion BT sCMOS cameras, was used for fluorescence microscopy of the hypotonic dissolution assay on the viral condensates. A Nikon CFI Plan Apo Lambda D 60x oil objective was used with 405 and 488 lasers for this study. 405 nm laser were used with 5% laser power and 50 ms exposure time to visualize Hoechst DNA stain, and 488 nm laser at 10% and 50 ms exposure to visualize reconstituted Ebola viral condensates labeled by mNG-VP35 in live HEK 293T cells. A Mad City Labs piezo z stage was used for z-stack acquisition. All acquisitions were performed with the Fusion BT in “Ultra Quiet” mode, and no binning was used.

The hypotonic dissolution assays were acquired via an automated imaging protocol created using the JOBS module of the Nikon NIS-Elements software. Just before imaging, the medium in each well in the 96-well plate was lowered from 200 µl to 50 µl of DMEM complete media. The automated imaging protocol included the following manual setup before imaging: 1) Define well selection and stage center position, 2) Set up the autofocus to scan 8 µm with 0.2 µm step size, 3) Designate the 488 nm and 405 nm lasers as the ‘Capture Definition,’ and 4) Pre-define a point with cells expressing wild-type or mutant viral condensates and with adequate Hoechst staining and assign its best z-plane for z position and perfect focus. After defining these parameters, the automated imaging was started, and sequential time lapse movies were collected once in each well according to the following order of events. 1) For the first 75 seconds, one image every 15 seconds, find autofocus using 405 nm laser at 5% laser and 10ms (same setting used for all autofocus in this protocol), then capture an image using the previously defined ‘Capture Definition’, 2) Wait and add 100 µl of water to the open top of the well to dilute culture medium from ∼150 mM to ∼50 mM as hypotonic treatment, or add 100 µl of complete medium in the control well as isotonic treatment, 3) For 15 minutes, acquire images with no delay time by performing autofocus, and capturing using ‘Capture Definition’, at ∼7.5 seconds per frame.

All image analysis for hypotonic treatment was performed in Fiji (ImageJ2)^104^ and Python v3.10. Files in .nd2 format were saved in two files, one capturing the pre-treatment and another capturing the 15 minutes immediately after water addition. These movies were concatenated using Fiji concatenate function to be used for analysis. Cell masks were generated in Python using Cellpose ‘cyto2’ model^95^ on mNG-VP35 and ‘nuclei’ model on Hoechst-stained nucleus to generate cytoplasmic and nuclear masks, respectively, using the first frame of the movie. Movies were then cropped per cell using the cytoplasmic masks as bounding boxes with a buffer of 10 pixels and saved as .tiff files. Bounding boxes were applied to all frames of the movie to obtain a cropped movie for each cell. The cropped movies were run through Rigid Body Registration using the nuclear channel to correct for bulk motion of cells for more accurate alignment with the masks over time. The registered cropped movies were then masked using the first frame masks of nucleus and cytoplasm, and cytoplasmic and whole cell mean intensities and standard deviations of intensities were measured at each frame using numpy.

The normalized coefficient of variation (CV) of mNG-VP35 mean intensity (per cell) was plotted over time using matplotlib. All time points were normalized to the average mean mNG-VP35 intensity of the first 5 pre-treatment frames. Error bars represent SEM across 4-5 wells per condition and mean calculated for isotonic wt, hypotonic wt, isotonic npssRNAmut, hypotonic npssRNAmut across n= 65, 115, 66, 75 cells, respectively. An exponential decay model was fitted based on the interval where the sharpest dissolution occurred (first 5 minutes), and shaded error is expressed as a 95% confidence interval.

### MERFISH sample preparation and imaging

#### Probe design and sample preparation

A total of 136 genes were identified and evaluated using the MERSCOPE^®^ Gene Panel Design Portal available at Vizgen (portal.vizgen.com) to ensure that each gene was suitable in length for probe binding and that abundance was below the recommended threshold to avoid optical crowding during imaging.

Human HEK293T cells were seeded on UV-sterilized MERSCOPE Slides (Vizgen, 20400001) placed in each 60 mm cell culture dish to reach 50% confluency the next day. Cells were transfected with plasmids expressing viral condensate proteins, treated with universal type 1 IFN as described in the previous section, and fixed with 4% Paraformaldehyde-fixed (PFA). Fixed slides were washed two times with 5 ml of 1x PBS and incubated with 70% ethanol at 4°C.

#### Antibody staining and probe hybridization

Cell boundary staining was performed using Vizgen’s Cell Boundary Staining Kit (Vizgen, 10400009) per the user guide for fresh- and fixed-frozen sample preparation. A fully detailed, step-by-step instruction on MERSCOPE sample preparation is available at https://vizgen.com/resources/fresh-and-fixed-frozen-tissue-sample-preparation/.

Briefly, samples were washed with 5 ml of 1x PBS then blocked for 1 hour at room temperature in Blocking Buffer C Premix (Vizgen 20300100) with RNase inhibitor (New England Biolabs, M0314L) added at 1:20 dilution. For primary antibody staining, samples were incubated for 1 hour at room temperature with mouse anti-Zaire Ebola VP35 (Kerafast, Kf-Ab01229-2.0) and human anti-Ebola NP nucleoprotein clone KZ51 monoclonal antibodies (Absolute Antibody, Ab00692-10.0), both at 1:100 dilution and RNase inhibitor (New England Biolabs M0314L) at 1:10 dilution in blocking buffer, followed by three washes with 5 ml of 1x PBS. Samples were then incubated for 1 hour at room temperature with Vizgen Protein Stain Anti-mouse and Anti-human Antibodies (Vizgen 20300101 and 20300105, respectively) at 1:100 dilution and RNase inhibitor (New England Biolabs, M0314L) at 1:10 dilution in blocking buffer, then washed three times with 5 ml of 1x PBS, postfixed in 5 ml 4% PFA-PBS for 15 minutes at room temperature, and washed twice with 5 ml of 1x PBS. Samples were then incubated in Formamide Wash Buffer (Vizgen 20300002) for 30 minutes at 37°C followed by probe hybridization with 50 µl of a custom designed MERSCOPE Gene Panel Mix at 37°C for 36 to 48 hours.

#### Gel embedding and clearing

Following incubation, the cells were washed twice with 5 ml of Formamide Wash Buffer (Vizgen 20300002) at 47°C for 30 minutes and embedded into a hydrogel using the Gel Embedding Premix (Vizgen 20300004), ammonium persulfate (Sigma, 09913-100G) and TEMED (N,N,N’,N’-tetramethylethylenediamine) (Sigma, T7024-25ML) from the MERSCOPE Sample Prep Kit (Vizgen 10400012). After incubation at room temperature for 1.5 hour, the samples were cleared with a solution consisting of 50 µl of Proteinase K (NEB, P8107S) and 5 ml of Clearing Premix (Vizgen 20300003) at 37°C for at least 24 hours, or until the cells became transparent.

#### MERFISH Imaging

After removing clearing solution and washing three times with Sample Prep Wash Buffer (Vizgen 20300001), the samples were stained with DAPI and Poly T Reagent (Vizgen 20300021) for 15 minutes at room temperature, washed for 10 minutes with 5 ml of Formamide Wash Buffer (Vizgen 20300002), then washed for 5 minutes with 5 ml of Sample Prep Wash Buffer (Vizgen 20300001). The imaging reagents and processed sample were loaded into the MERSCOPE and a low-resolution DAPI mosaic (10X magnification) was used to select the region of interest before high-resolution imaging on the MERSCOPE system (Vizgen 10000001). With a 60X oil immersion objective, DAPI and Poly T stains were imaged at 7 focal planes on the z-axis for each tiled field of view (FOV), followed by 6 rounds of 3-color imaging across all focal planes using 750 nm, 650 nm, and 560 nm laser illumination. Each imaging round was followed by incubation in extinguishing, rinse, hybridization, wash, and imaging buffers. Additionally, a single image of fiducial beads was acquired at each FOV using 488 nm illumination. The full instrumentation protocol is available at https://vizgen.com/wp-content/uploads/2021/07/91600001_MERSCOPE_Instrument_User_Guide_RevJ.pdf.

#### MERFISH pre-analysis transcript detection

The MERlin image analysis pipeline^105^ was used to analyze the raw image files from the MERSCOPE experiment to align image stacks from the different MERFISH rounds, filter out background noise, and enhance RNA spot detection. Individual RNA molecule barcodes were then decoded using a pixel-based algorithm and an adaptive barcoding scheme that corrects misidentified barcodes not matching the provided codebook.

### MERFISH subcellular analysis

#### Cell and Compartment Segmentation

All MERSCOPE outputs (multi-channel TIFF stacks and positional mapping files) were first converted into Zarr^106^ format via the merscope function from scverse’s spatialdata-io package^107^. Zarr arrays and metadata were loaded into a SpatialData object for unified handling of image channels and spatial coordinates. Three fluorescence channels, DAPI, Poly-T, and VP35, were preprocessed for downstream segmentation. For the DAPI channel, a morphological erosion (5×5 kernel) using OpenCV^108^ was applied to tighten the stain boundaries and compensate for intensity blurring. The Poly-T channel underwent the same erosion step followed by local histogram equalization using the scikit-image^109^ equalize function to balance nuclear and cytoplasmic intensities and improve cell-boundary detection in downstream cell segmentation. VP35 images were processed with the scikit-image equalize adapthist function for contrast-limited adaptive histogram equalization (CLAHE) to amplify condensate contrast, then smoothed with the scikit-image Gaussian filter to reduce noise and soften edges.

Cell and nuclei segmentation tasks were executed in parallel using Sopa^110^, which subdivided each image into 5000×5000-pixel tiles with 200-pixel overlaps, processed each tile independently, and then merged the results into seamless, contiguous segmentation masks. Nuclear masks were generated by the Cellpose^95^ *nuclei* model on the DAPI channel preprocessed with erosion. Whole-cell segmentation employed the Cellpose *cyto3* model on the preprocessed DAPI and Poly-T channels. Nuclear masks were generated by the Cellpose *nuclei* model on the preprocessed DAPI channel. Condensates were segmented by multi-Otsu thresholding of the VP35 channel, extracting the highest-intensity classes as candidate condensate objects. Condensate masks were refined using the Bento^96^ segmentation overlay function for compartment delineation and filtering, and the shape analysis function for morphometric calculations. Segmentation overlay discarded objects overlapping nuclear regions or extending beyond cell boundaries. Morphometric filtering then removed condensates smaller than 20 pixels or larger than 50% of their host cell’s area. A final cytoplasmic compartment mask was created by subtracting the refined condensate regions from the total cytoplasmic compartment.

#### Transcript Localization Evaluation and Thresholding

Single-molecule transcript coordinates from MERFISH^111^ were assigned to nuclear, cytoplasmic, or condensate compartments via point-in-polygon tests using the Shapely^112^ and Geopandas^113^ libraries. For each gene-derived transcript, we calculated the Log_₂_ (condensate-to-cytoplasm transcript density ratio), defined as the log2 transformed mean transcript density within condensates divided by that in the cytoplasm, to quantify relative enrichment of transcript in viral condensates. To derive transcript density ratio, we divided the total transcript count in each compartment by the compartment area. Transcript localization metrics include the %cell with detected transcript (defined as the proportion of cells containing at least one transcript of a specific gene), the mean transcript count per cell, and compartment specific measurements (compartment area by pixel^2^, total transcript count per compartment, transcript density, and compartment transcript density normalized to that of the whole cell).

Cells without detected condensates and cells with minimum total transcript detected (below 5% of all cells measured) were excluded from the calculation of transcript localization metrics. All MERFISH-detected transcripts were thresholded by excluding transcripts with mean count lower than the highest mean count of misidentified false positive control (blank control^48^). Detected transcripts passing this threshold were considered as high-confidence hits. Within high-confidence hits, a subset of transcripts with mean count per cell >3 and condensate-to-cytoplasm transcript density ratio >1 were further designated as “condensate-enriched”. Transcripts belong to this subset, have on average more than 3 copies per cell that allows meaningful partitioning into more than one subcellular compartment and have higher concentration of transcripts in the viral condensates than in cytoplasm. Raw data of MERFISH subcellular analysis are shown in Table S1.

### RNA folding prediction and feature analyses

High-confidence, condensate-enriched transcripts in each experimental condition (-/+ type 1 IFN) were subjected to sequence and structural feature analyses. Transcript length and GC content were extracted from reference FASTA sequences. Full-length transcripts were analyzed for the predicted number of G-quadruplex within the input sequence using QGRS Mapper^102^. RNA secondary structure predictions were generated using RNAfold from the ViennaRNA Package^54^, computing thermodynamic ensemble free energies for full-length transcripts, coding sequences, 5′ UTRs, and 3′ UTRs. Importantly, thermodynamic ensemble free energies, which account for the heterogeneity of intracellular RNA folding^114^, were used instead of the minimum free energies for input sequence. If not specified, all RNA folding predictions use physiological temperature (37°C) and salt concentration (150 mM) as energy parameters. These energy values (kcal/mol) as well as the predicted number of G-quadruplex were divided by sequence length (nt) to adjust for length-dependent effect. The transcript length, GC content, length-normalized G-quadruplex number and length-normalized RNA folding were correlated with the log transformed, condensate-to-cytoplasm transcript density ratios for each transcript using Spearman’s correlation and simple linear regression. Spearman’s r value and p values were reported for all features being analyzed. The R^2^ and P values (*, P<0.05, **, P<0.01, ***, P<0.001) from linear regression were reported for features with significant Spearman’s correlation.

For condensate-enriched, high-confidence hits from the interferon-treated dataset, additional RNA folding analyses were performed for orthologous transcripts from Egyptian fruit bat (*Rousettus aegyptiacus)*, domestic ferret (*Mustela putorius furo)*, and rhesus macaque (*Macaca mulatta*) using 37°C and 39°C as energy parameter. The length-adjusted RNA folding free energy for selected human genes were correlated with the non-human orthologous using simple linear regression. Slope and R^2^ values were reported for those slopes that are significantly non-zero.

For negative-strand RNA viruses, the full-length, sometimes multicistronic RNA genome contains not only the reverse complement sequence of each protein coding gene, but also the 3’leader and 5’ trailer sequences, intergenic sequences, and the noncoding regions (reverse complement of UTRs) of each gene cassette. The motivation of analyzing viral sequences is based on the strong correlation between the RNA folding of cellular mRNA coding sequence and viral condensate enrichment. To make a fair comparison in the RNA folding of viral and cellular RNA coding sequence, the coding sequence of individual viral genes belonging to each virus were used.

All sequences used in RNA folding analysis are shown in Table S1.

### RNA-IP and RT-qPCR

RNA-IP cell lysate was prepared based on previous studies^115–117^. HEK 293T cells were seeded at 6X10^6^ cells per 10 cm^2^ dish 24 hours prior to transfection. Each dish of cells with 7 ug of pCAGGS-NP plus 7 ug of pCAGGS-HA-mNG-VP35 plasmids using Trans-IT LT1 transfection reagent. pCAGGS-VP35 without epitope tag was used to replace tagged VP35 in the control condition. At 24 hours post transfection, prepare cell lysate (>=10^7^ cells) in cold, autoclaved hypotonic gentle lysis buffer (10 mM Tris pH7.4, 10 mM NaCl, 10 mM EDTA, 0.5% Triton X-100, freshly supplied with Halt^TM^ Protease inhibitor cocktail and phosphatase inhibitor cocktail (Thermofisher, 78445)) using a 21 G needle connected with sterile syringe. Lysates were clarified at 12,500 xg for 10 minutes. In clarified lysates, autoclaved NaCl stock was added to reach a final concentration of 150 mM and nucleases-free glycerol stock was added to a final concentration of 5% (v/v). Clarified lysates were flash frozen in liquid nitrogen and stored at – 80 °C freezers for later use, if not proceed to protein quantification and RNA-IP the same day.

Total protein quantity in each clarified lysate sample was measured using BCA assay and samples containing 5 mg of total proteins were used for immunoprecipitation with Roche anti-HA affinity matrix (Sigma, 11815016001) (100 µl of beads volume per 5 mg sample). 0.5% of the lysate sample was saved for protein input in western blotting, 2.5% was saved for RNA input in RT-qPCR. Anti-HA beads were blocked with autoclaved NT2-buffer (50 mM Tris-HCl (pH 7.4), 150 mM NaCl, 1 mM MgCl2, and 0.05% NP-40) with 0.5 mg/ml of invitrogen ultrapure BSA (Thermofisher, AM2616). Blocking took place in a rotator for at least 30 minutes at 4°C. Blocked anti-HA beads were combined with lysate samples containing equal total proteins and incubated in a rotator for at least 2 hours at 4C. Beads were spun down at 10,000 xg for 30 seconds and unbound fractions were collected for western blotting (0.5% of total supernatant) and RT-qPCR (2.5% of total supernatant). Beads were washed 4 times with 10 beads-volume each of the autoclaved NT2 buffer. Wash beads were separated into two parts, 2% of total beads was boiled in NuPage^TM^ LDS sample buffer (4X) (Thermofisher, NP0007) for western blotting, the rest of beads reconstituted in Trizol LS for total RNA extraction using Zymo Direct-zol RNA miniprep kit (ZymoResearch, R2050).

RNA quality in each sample was evaluated by loading < 10 ng of total RNA/sample in high sensitivity RNA screentape for the presence of 18s and 28s ribosomal RNAs via Agilent TapeStation analysis. Equal volume of nuclease-free water was used to elute input/unbound/IP fractions across all samples and equal volume of input/IP-derived total RNA was used for cDNA synthesis reaction using the High-capacity cDNA Reverse Transcription kit (Thermofisher, 4368814) paired with the Oligo(dT)_20_ primer (Thermofisher, 18418020). Reverse transcription of cDNA was performed following the manufacturer’s protocol. Equal volume of cDNA in each sample was diluted, mixed with target specific qPCR primer pairs from PrimerBank^118^, and with Applied Biosystems™PowerUp™ SYBR™ green master mix for qPCR (Thermofisher, A25742). The relative quantification (ΔΔCq) method was used to analyze qPCR amplification results. Sequences of all qPCR primers are shown in Table S3.

### Infection with biologically contained Ebola virus

HEK293T cells were seeded in 6-well plate (2X10^5^ cells/well) and transfected with pCAGGS-Ebola VP30 (3.75 ug of plasmid/well) the day after using lipofectamine 3000. One-day post transfection, transfected HEK293T cells (4X10^4^ cells/well) were reseeded in human fibronectin coated glass-bottom IBIDI 8-well u-slide. Twenty-four hours later, the monolayer was incubated with EBOV-ΔVP30-GFP^64^ at MOI=5. After 6 hours of incubation at 37°C, the monolayer was washed three times with DPBS to remove extracellular virions and instead incubated with DMEM containing 2% FBS and recombinant universal type 1 IFN at 1000U/ml for 18 hours. Infection was terminated at 24 hours. Infected cells were inactivated with 4% PFA-PBS for 15 mins. Fixed samples were used for SABER-FISH and immunofluorescence analysis.

To quantify the size of viral condensates and partition coefficient of endogenous transcript ODC1 and IFIT3 in infected cells, an image analysis pipeline was built in CellProfiler 4.2.6^103^. Immunofluorescence labeled NP channel was used for viral condensate segmentation, Hoechst channel for nuclei segmentation, and Gaussian-filtered (Sigma=10) FISH channel combined with diffused GFP signal (expressed by the reporter virus) for cytoplasm segmentation. Cytoplasm segmentation was performed using Cellpose 2.3.2^95^ module under cyto2 detection mode. Viral condensate segmentation was performed using IdentifyPrimaryObjects module with Two classes, Global thresholding, and Otsu. FISH-labeled transcript detection and quantification is described in section below.

### Design and expression of synthetic reporter mRNAs in cells

To construct the synthetic reporter with mScarlet coding variant, termed pCAGGS-mScar-CDS reporter, two synonymous mScarlet coding sequences were generated by replacing GC-rich codons with AT-rich codons using the manual optimization option under IDT Codon Optimization Tool and synthesized as gene fragment by IDT and Genscript. The human ODC1-5’ and 3’UTR fragments were obtained from PCR amplification using cDNA of whole cell RNA isolated from HEK293T cells as the template. These three DNA fragments were assembled into the pCAGGS backbone using NEB HiFi Assembly kit.

To construct the IFN-inducible synthetic reporter with varying 3’UTRs, termed pCDNA3-ISRE-mScar-3’UTR reporter, four DNA fragments were assembled into the pCDNA3 plasmid backbone, omitting its original CMV enhancer and CMV promoter. These fragments include IDT synthesized gblock encoding the -300 to transcription start site (TSS) of human ISG15 promoter sequence that contain ISRE, human OASL 5’UTR sequence amplified from cDNA of IFN-stimulated HEK293T cells total RNA, mScarlet Mod1 coding sequence, human SAMHD1/IFIT1/IFI6 3’UTR amplified from cDNA of IFN-stimulated HEK293T cells total RNA, or Egyptian fruit bat SAMHD1/IFIT1/IFI6 3’UTR cDNA fragment synthesized by IDT. Sequence of positive clones confirmed by plasmidsaurus sequencing.

To express synthetic reporter with mScarlet coding variant for RNA-FISH analysis, HEK293T cells seeded in IBIDI 8-well u-slide were transfected with a combination of 50 ng/well of pCAGGS-NP, 50 ng/well pCAGGS-HA-mNG-VP35, and 10 ng/well of each pCAGGS-mScar-CDS reporter plasmid using Trans IT LT-1 transfection reagent.

To express IFN-inducible synthetic reporter with varying 3’UTR for RNA-FISH analysis, HEK293T cells seeded in IBIDI 8-well u-slide were transfected with a combination of 50 ng/well of pCAGGS-NP, 50 ng/well pCAGGS-HA-mNG-VP35, and 50 ng/well of each pCDNA3-ISRE-mScar-3’UTR reporter plasmid using Trans IT LT-1 transfection reagent. Transfected cells were stimulated with universal type 1 IFN at 1000 U/ml at 5-hours post transfection. Annotated sequence of all reporters is summarized in Table S4.

### SABER-FISH sample preparation, imaging, and image quantification

HEK293T (3-4X10^4^ cells/well) were seeded in human fibronectin coated glass-bottom IBIDI 8-well u-slide 24 hours prior to transfection. Cells were co-transfected with 50 ng/well of pCAGGS-NP or pCAGGS-NPssRNAmut paired with 50 ng/well pCAGGS-HA-mNG-VP35 using Trans IT LT-1 transfection reagent. For IFN-stimulated samples, transfected cells were incubated with complete DMEM-containing 1000U/ml recombinant universal type 1 IFN about 4-5 hours post transfection for 24 hours stimulation. To reconstitute Ebola virus minigenome RNA replication, cells were co-transfected with previously described^35^ minigenome system: pCAGGS-NP (25 ng/well), pCAGGS-VP35 (25 ng/well), pCAGGS-VP30 (15 ng/well), pCAGGS-T7pol (100 ng/well), pCAGGS-L (200 ng/well), and pT7-2cis-MG-replication competent that encode both eGFP and Renilla luciferase (100 ng/well) using Trans IT LT-1 transfection reagent. At 1 day post transfection, samples were chemically fixed with 4%PFA in PBS for 10 minutes, washed 3x 5 minutes in 1x PBST and proceeded to the SABER FISH protocol as previously described^119^.

Briefly, samples were hybridized with 1 µg of concatemerized pooled probes targeting ODC1 mRNA or IFIT3 mRNA alone or combined with 1 µg of concatemerized probe targeting Ebola virus minigenome reporter vRNA per well for 16 hours at 43 °C. Pooled probes containing 14 distinct DNA oligos complementary to the coding sequence of human ODC1 were concatemerized with hairpin 27 in a Primer Exchange Reaction (PER) with 1 hour extension time according to published protocol^52^.

A single probe, EBOV_tr_25p, targeting the 5’ trailer sequence in the Ebola virus genome was concatemerized with hairpin 25 using 1 hour extension time in PER. Sample incubation with fluorescence imager oligos (0.4 µM each probe, for imaging probes with hairpin 25: /5ATTO565N/tt TATTATTGGT TATTATTGGT /3InvdT/; for probes concatemerized with hairpin 27: /5ATTO647NN/tt ATGATGATGT ATGATGATGT /3InvdT/) was performed at 37°C for 10 minutes. For subsequent immunofluorescence staining, primary and secondary antibodies were diluted in Nuclease-free 1x PBST for desired period with washing step (3x 5 minutes in 1x PBST) following each incubation.

To detect synthetic mScarlet reporter mRNAs, pooled probes containing 14 distinct DNA oligos complementary to the coding sequence of each variation of mScarlet coding sequences and were concatemerized with hairpin 27. All primary FISH probe sequences are manually designed to have 20-30 nucleotides annealed to target sequence, Tm between 60 to 70°C, and GC content between 40-60%, and confirmed to have no predicted binding to other human transcripts expressed in the tested cell line using BLASTn analysis. Sequences of primary FISH probes are listed in Table S2.

To quantify endogenous ODC1 mRNA detected by SABER-FISH, an IdentifyPrimaryObject module was added to the previously mentioned CellProfiler pipeline, in which a constant diameter range (Min,Max) was used to identify the mRNA puncta as individual primary object. Immunofluorescence-stained XRN1 channel was used to segment cytoplasm, Hoechst channel was used to segment nuclei, and mNG-VP35 channel was used to segment viral condensates. Area of identified cytoplasm and viral condensates was measured using the MeasureObjectSizehape module. mRNA puncta count was normalized to the measured area of either cytoplasm or viral condensates and was compared across condensates containing wild type NP or RNA binding mutant. For quantification of endogenous IFIT3 mRNA, due to the variation of IFIT3 mRNA signals within and across individual cells (because of heterogeneous IFN response), the mean intensity of FISH-labeled mRNA signal in the compartments of interest was used for partition coefficient measurement.

To quantify mScarlet reporter mRNA detected by SABER-FISH, a similar CellProfiler image analysis pipeline was used with the following modifications. Due to the high expression level of mScarlet reporters in transient transfection experiments, instead of detecting individual mRNA puncta, the mean intensity value of FISH-labeled mRNA channel in viral condensates and in non-condensates area in cytoplasm were used to calculate the partition coefficient of each reporter mRNA in viral condensates. A combination of FISH-labeled mScarlet mRNA channel and mScarlet fluorescence was used to segment cytoplasm, Hoechst channel was used to segment nuclei, and mNG-VP35 channel was used to segment viral condensates.

### XRN1 and EIF4G immunofluorescence analysis

HEK293T cells were seeded, transfected, in which endogenous ODC1 mRNAs were stained as described in the SABER-FISH section. After completion of SABER-FISH sample preparation, samples were further incubated with either anti-eIF4G mouse monoclonal antibody (A-10) (sc-133155) at 1: 100 or anti-XRN1 mouse monoclonal antibody (C-1) (sc-165985) at 1:500 for 4 hours at room temperature, washed with RNase-free PBS for three times, and incubated with goat anti-mouse Alexafluor568 (1:1000) for 1 hour, washed again with RNase-free PBS for three times. Nuclei were stained with Hoechst for 15 minutes, after which samples were mounted in ProLong^TM^ Gold anti-fade mountant and the next day imaged on Zeiss LSM980-airyscan system using airyscan-MULTI mode.

To quantify XRN1 and EIF4G partition, the CellProfiler image pipeline described above was modified to use XRN1 or EIF4G immunofluorescence channel for cytoplasm segmentation. Due to the granular staining pattern of both XRN1 and EIF4G, upper quartile intensities for each protein in viral condensates or in cytoplasm were used for partition coefficient quantification.

### Locked nucleic acid (LNA)-antisense oligo (ASO) design and treatment

A 17-mer LNA/DNA mixer ASO targeting a predicted single-stranded region in human ODC1 coding sequence was designed manually to meet the following criteria: 1) GC-content between 30-60%, 2) no stretches of 3 or more Gs or Cs, 3) discontinuous LNA modification to break stretches of 4 or more unmodified DNA, 4) RNA Tm above 80°C calculated by the Qiagen LNA Tm prediction online tool, 5) the hybridization score and secondary structure score below 30°C calculated by the Qiagen LNA oligo optimizer online tool, 6) no off-target binding to other human transcripts expressed in the tested cell line using BLASTn analysis. Two phosphorothioate (PS) bonds were added to both 5’ and 3’ end of the ASO to improve stability and block RNaseH cleavage. Modified ASO was synthesized by IDT with HPLC purification.

To sterically block endogenous ODC1 mRNA recruitment into viral condensate, HEK293T cells transiently transfected with Ebola NP and VP35 expression plasmids were transfected with the ODC1 targeting LNA-ASO 5 hours after the first transfection, using Lipofectamine 3000.

### Western blotting and RT-qPCR analysis on condensate-enriched target

HEK293T cells (1×10^6^ cells/ well) were seeded in 24 wells plate and co-transfected the next day with pCAGGS-NP wild-type and pCAGGS-mNG-VP35 wild-type at 300 ng or 600 ng per plasmid per well using Trans-IT LT1 transfection reagent. Twenty-four hours post transfection; whole cells lysate or total RNA was collected. To collect whole cell lysates, cell pellets pooled from three replicate wells were first resuspended in 1xRIPA buffer (Cell signaling, #9806) supplied with Halt^TM^ proteases and phosphatase inhibitor cocktail and Benzonase nuclease, ultrapure (Sigma, E8263) then clarified into soluble lysates at 12,000 xg for 5 minutes. Clarified lysates were mixed with reducing agent and sample loading buffer, boiled at 95 °C for 5 mins, then loaded in NuPAGE^TM^ Bis-Tris Mini Protein Gels, 4-12%, (1.5 mm, 10-well) (Thermofisher, NP0335BOX) and transferred to PVDF membrane.

To probe for target proteins, membrane was sequentially incubated with primary antibodies targeting GAPDH (1:4000 dilution, SantaCruz, sc-32233), DCP1A (1:2000 dilution, Abcam, ab183709), ODC1 (1:1000 dilution, Proteintech, 28728-1-AP), EIF2AK2/PKR (1:1000 dilution, Abcam, ab184237), IFIT3(1:1000 dilution, Sigma, HPA059914), RIG-I (1:1000 dilution, Thermofisher, MA5-31715), and RSAD2 (1:1000 dilution, Proteintech, 28089-1-AP) diluted in the blocking buffer (5% BSA-TBST) for either 2 hours at room temperature or overnight at 4°C, followed by 3x TBST washes. Secondary antibodies, Peroxidase AffiniPure™Goat anti-Rabbit IgG (H+L) (Jackson ImmunoResearch, 111-035-144) or anti-Mouse IgG (H+L) (Jackson ImmunoResearch, 115-035-062), were 1:2000 diluted in the blocking buffer and incubated with membrane for 1 hour at room temperature, followed by 3x TBST washes. Clarity western ECL substrate (BioRad, 1705061) was incubated with membrane to produce chemiluminescent signal, which was imaged by a BioRad Gel Doc system and quantified in Fiji (ImageJ2). In Fiji (ImageJ2), each lane was manually selected via Analyze>Gels>Select First/Next Lane and plotted by the Plot Lanes function. Intensity of the target protein in each lane was calculated by measuring areas under the curve (AUC) using the wand tool.

Total RNA from each sample pooled from three replicate wells were extracted using Zymo Direct-zol RNA miniprep kit (ZymoResearch, R2050). Samples from each condition containing 200 ng of total RNA were used for cDNA synthesis. Equal volume of diluted cDNA samples used in qPCR for relative quantification of target mRNA compared to the reference mRNA as described in the previous section. The sequence of all primers used in qPCR are provided in Table S3.

### Cellular mRNA turnover analysis with Actinomycin D treated cells

HEK293T cells (1×10^5^ cells/ well) were seeded in 24 wells plate and co-transfected the next day with pCAGGS-mNG-VP35 wild-type plus pCAGGS-NP wild-type or NP-ssRNA binding mutant at 600 ng per plasmid per well using trans-IT LT1 transfection reagent. At twenty-four hours post transfection, cells were treated with Actinomycin D at 5 ug/ml to block translation. Whole cell RNA was collected in Trizol at 0, 1, 2, 4, 8, 24 hours post Actinomycin D (Sigma, A4262) incubation and extracted using Zymo Direct-zol RNA miniprep kit (ZymoResearch, R2050). Duplicate wells in each time point for each condition were pooled into one sample. Samples from each time point containing 200 ng of total RNA were used for cDNA synthesis. Equal volume of diluted cDNA samples used in qPCR for relative quantification of target mRNA compared to the reference mRNA as described in the previous section.

### Quantification and Statistical analysis

All statistical analyses were performed in GraphPad Prism 10. Statistical details of experiments, including the type of test used, values and error bars displayed, the number of independent experiments/replicates, can be found in all figure legends.

In summary, Mann-Whitney test was used to determine the difference between independent conditions in microscopy experiments, given small sample size and presence of outliers in single cell measurements. Paired t-test was used to determine difference between compartment-specific transcript density in MERFISH data, given large sample size and the paired relationship between condensates and cytoplasm in each cell analyzed. Paired t-test was also used to determine difference in mRNA/protein levels in the presence to absence of viral condensates in four independent experiments, data points across different conditions in the same experiment were paired. Unpaired t tests with Welch’s correction were used to determine the IFN-induced ISG mRNA upregulation in one representative experiment with technical triplicates and to determine the RNA folding difference between selected cellular mRNAs and representative viral RNAs. Spearman’s correlation and simple linear regression were used to determine the correlation between RNA features and condensate-enrichment. Simple linear regression was used to determine the evolutionary conservation in RNA folding between human and non-human mammals for a small selection of genes, with the slope value indicating one species has stronger/weaker RNA folding compared to the other species.

## Supplemental video and Excel table titles and legends

**Video S1. Time lapse of a live HEK293T cell co-expressing mNG-VP35 plus NP-wt before and after hypotonic shock.** Elapsed time, 20 min. Scale bar, 10 μm. Related to Figure S2.

**Video S2. Time lapse of a live HEK293T cell co-expressing mNG-VP35 plus NP-ssRNA mut before and after hypotonic shock**. Elapsed time, 20 min. Scale bar, 10 μm. Related to Figure S2.

**Table S1. Cellular mRNA and viral RNA sequence and corresponding RNA folding free energy predicted by RNAfold, MERFISH subcellular quantification metrics, species-specific RNA folding comparison.** Related to Figure 2B, 2D, 2F, Figure 4B, 4C, 4E, 4G, Figure S3E, Figure S4C

**Table S2. Sequences of primary FISH probes used in SABER-FISH based detection of endogenous mRNAs, viral RNA genome, and reporter mRNAs.** Related to the Key Resource Table in the STAR Methods

**Table S3. Sequences of all primers used in qPCR.** Related to the Key Resource Table in the STAR Methods

**Table S4. Sequences used in constructing all synthetic RNA reporters.** Related to Figure 2 and Figure 4

