## Supplemental Information for "RNA folding drives species-dependent host mRNA partitioning in Ebola virus condensates"

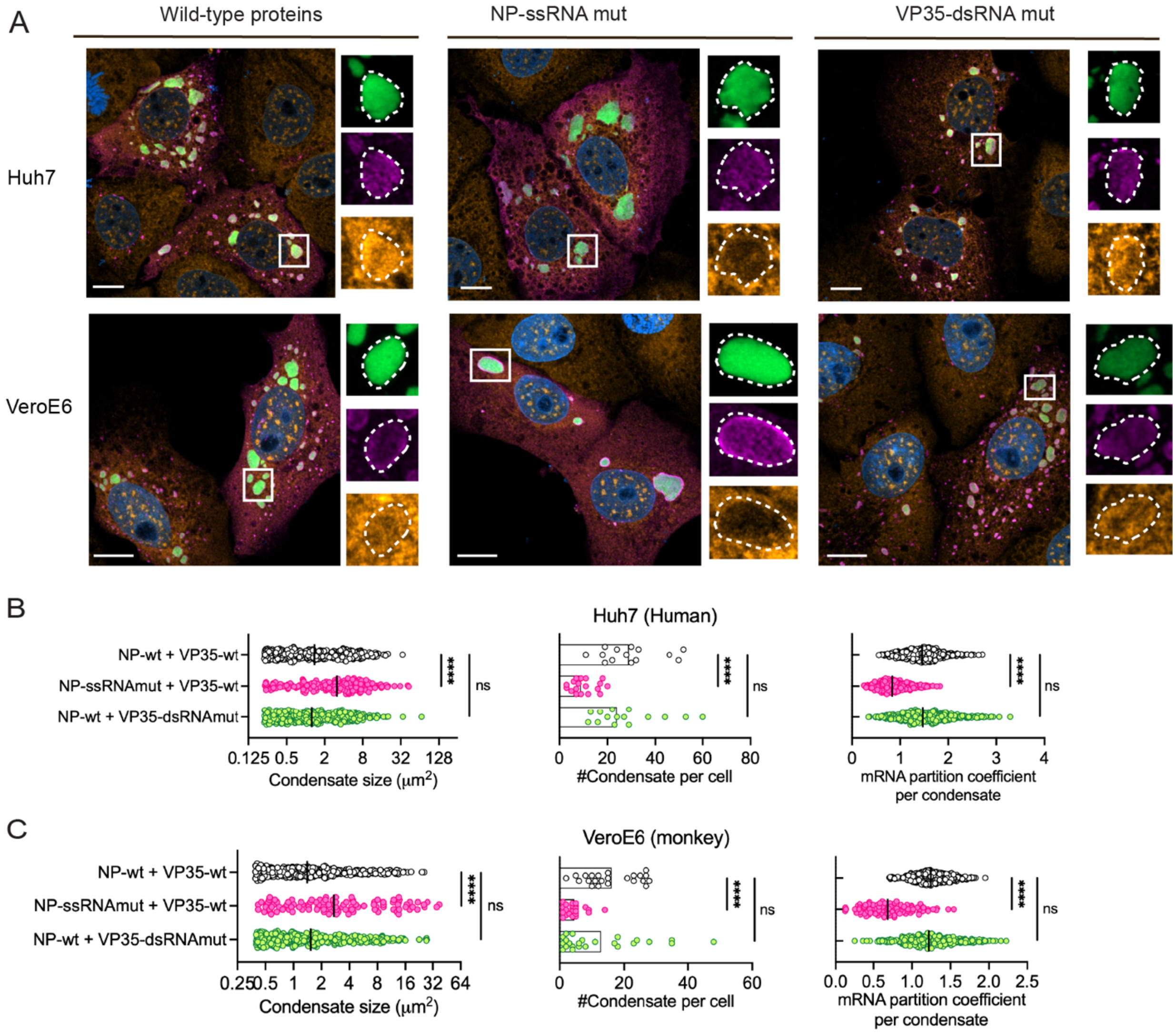


**Figure S1. Ebola virus condensates partition cellular mRNA in human liver and monkey kidney cells, related to Figure 1**

**A.** Confocal microscopy of fixed and stained Huh7 (human liver) and VeroE6 (monkey kidney) cells co-expressing untagged Ebola virus NP and mNG-tagged Ebola virus VP35 in either wild-type or RNA-binding mutant forms. Individual channels are shown for a representative condensate in each condition. Scale bar: 10 μm. Nuclei stained with Hoechst.

**B.** **C.** Condensate size, number and cellular mRNA partition coefficient for Ebola virus NP-VP35 condensates with either wild-type or RNA-binding mutant NP scaffold in Huh7 (B) and VeroE6 cells (C). Median values with individual data points (per condensate) from two experiments (with > 15 cells in each condition combined for

Huh7, >30 cells for VeroE6 are shown). ****, P<0.0001; ns: not significant, by Mann-Whitney test.

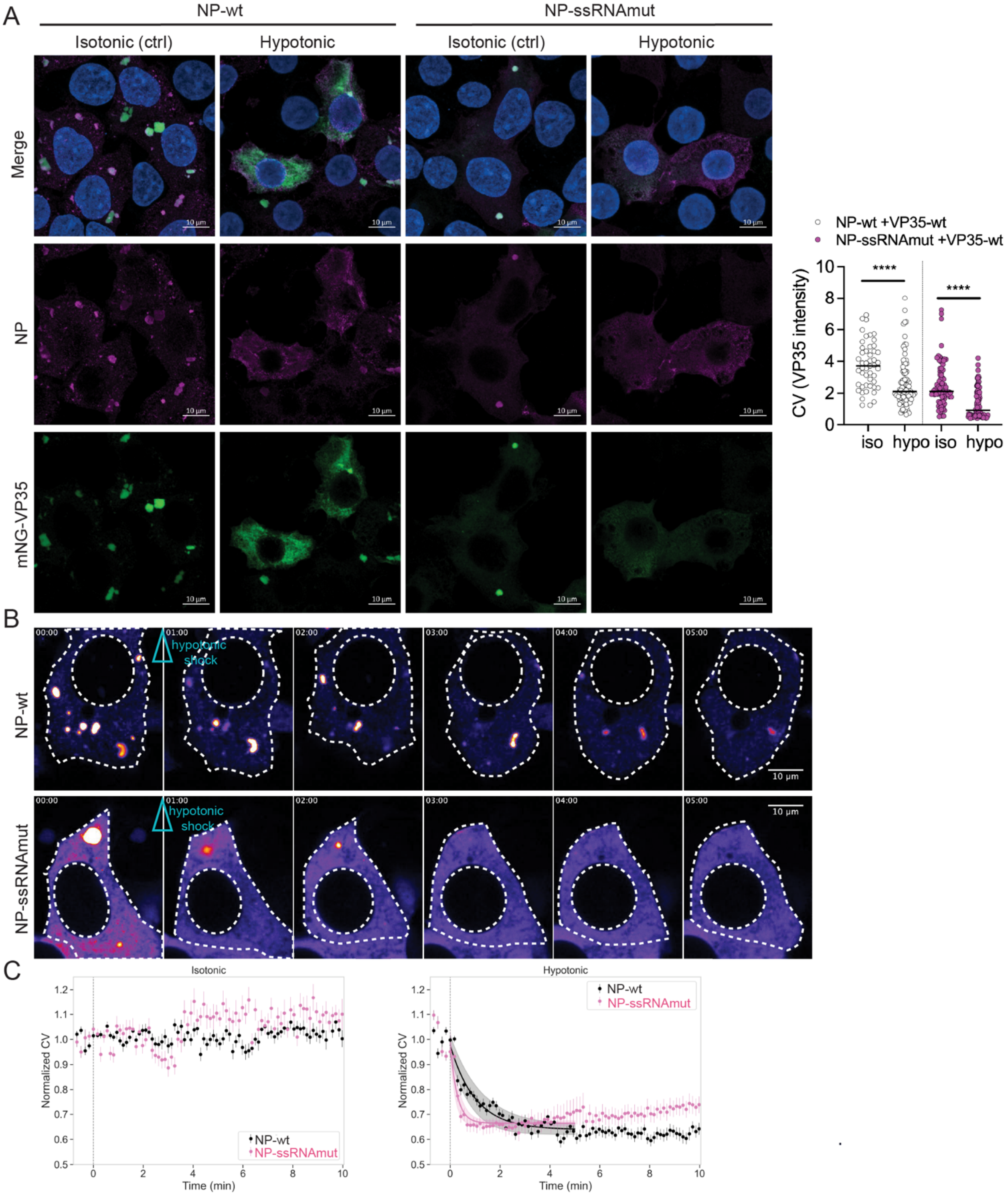

**Figure S2. RNA-containing and RNA-depleted viral condensates display different degrees of disassembly upon hypotonic treatment in living cells, related to Figure 1**

**A.** Confocal microscopy of fixed and stained HEK293T cells co-expressing mNG-tagged Ebola virus VP35 plus either Ebola virus wild-type NP (wt) or RNA binding mutant NP (ssRNAmut). Prior to fixation, live cells were incubated with either complete media (isotonic) or media diluted 2 times with water (hypotonic) for 15 minutes. Coefficients of variation (CV) of mNG-VP35 intensity per cell measured for either wild-type or mutant condensates under different conditions are shown. Representative results from two experiments are shown and quantified (>49 cells in each condition). ****, P<0.0001, by Mann-Whitney test.

**B.** Time lapse of live HEK293T cells co-expressing mNG-VP35 plus either NP-wt or NP-ssRNAmut before and after hypotonic shock. The mNG-VP35 channel at selected frames in the first 5 minutes are shown with the timing of hypotonic shock indicated by an empty arrowhead. Results of two experiments with >60 cells in each condition were quantified and plotted. CVs of mNG-VP35 intensity per cell in each frame after isotonic/hypotonic treatment were normalized to the average intensity of the first 5 frames prior to treatment. An exponential decay model was fitted for normalized CV values (Mean±SEM) in the first 5 minutes for hypotonic treated conditions. Shaded error is expressed as a 95% confidence interval.

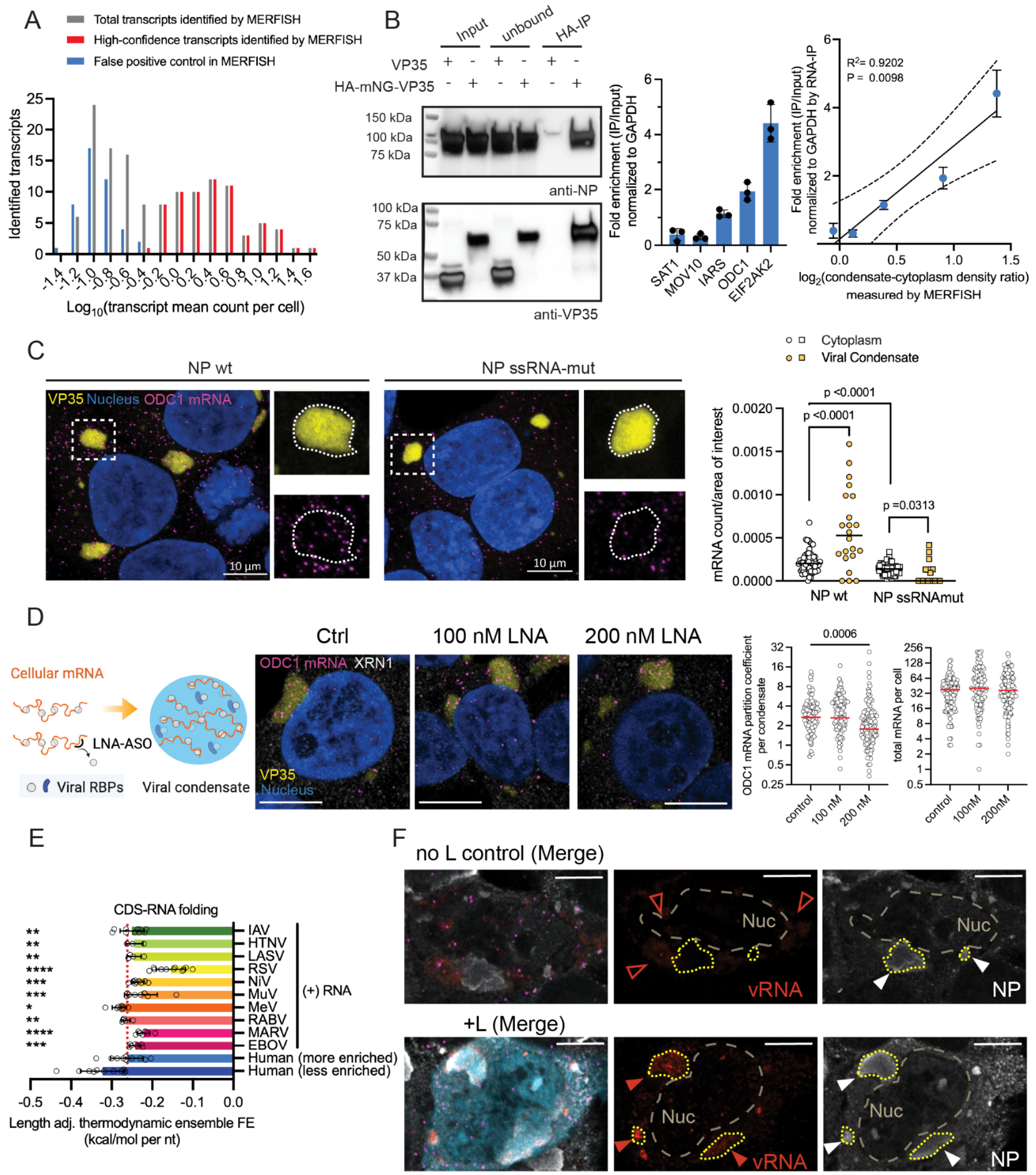

**Figure S3. MERFISH transcripts thresholding and validation, related to Figure 2**

**See also Table S1**

**A.** Distribution of MERFISH-identified transcripts ranked by mean count per cell for individual transcripts including 136 target genes (true positive) and 44 misidentification controls (false positive). The highest mean count value for control detected by MERFISH was taken as the cutoff for high-confidence transcripts.

**B.** RNA-coimmunoprecipitation (RNA-IP) validation of 5 representative MERFISH-detected transcripts sampling different degrees of condensate-enrichment. HEK293T cells co-transfected with Ebola NP plus untagged or HA-mNeonGreen (mNG)-tagged VP35 were lysed and soluble fraction containing equal total proteins used for HA-IP under RNase-free conditions. Anti-Ebola NP antibody and anti-Ebola VP35 antibody were used to detect NP in cell lysate input, unbound and HA-IPed fraction. The relative abundance of five representative targets plus GAPDH was quantified by OligodT-RT-qPCR using total RNA extracted from Input and IP fraction. GAPDH mRNA level in both fractions used as an arbitrary internal control for normalization. For each target, mean±SD of relative fold enrichment (IP-to-Input) is shown and further correlated with the condensate enrichment metric (condensate-cytoplasm density ratio) measured by MERFISH. Representative results of two experiments are shown. $R^2$, P value, and fitted lines with 95% confidence intervals from linear regression are shown.

**C.** Confocal microscopy of fixed and stained HEK293T cells co-expressing mNG-VP35 plus either wild-type Ebola NP (wt) or RNA-binding mutant NP (NP ssRNA-mut). Boxed condensate within a merge overview is magnified to show individual channels: condensate marker, mNG-VP35, and endogenous ODC1 mRNA puncta labeled by SABER-FISH. Cytoplasm was visualized by immunofluorescence staining of XRN1 and shown in white in the merge overview. Nuclei stained with Hoechst and shown in blue. Scale bar: 10 µm. Relative ODC1 mRNA density depicted by ODC1 mRNA puncta count normalized to area of interest are shown for both cytoplasm and viral condensates for both wild-type and mutant condensates. Representative results of three experiments (>10 cells in each condition per experiment) are shown. Mann-Whitney test used to determine the difference in ODC1 mRNA density in viral condensates versus the cytoplasm of the same cell. P values are shown.

**D.** Validation of NP-mediated ssRNA binding mediates ODC1 mRNA enrichment in viral condensate using antisense oligonucleotides (LNA-ASO) targeting ODC1 ssRNA region. Confocal microscopy of fixed and stained HEK293T cells co-expressing mNG-VP35 and wild-type Ebola NP and transfected with control (transfection reagent only) or ODC1 targeting LNA-ASO 100 at nM/200 nM. Scale bar: 10 µm. Representative results of two experiments (>120 cells in total per condition) are shown. One-way ANOVA used to evaluate the effect of LNA-ASO treatment on ODC1 mRNA partitioning and Kruskal-Walli test (multiple comparisons) used to compare the difference between the LNA-ASO-treated group to the control.

**E.** Comparison of the predicted RNA-folding energy (length-adjusted) for cellular and positive-strand virus CDS regions. Human (more/less enriched): high-confidence cellular mRNAs detected in MERFISH with top/bottom 50% ranked transcript enrichment in the viral condensates. HTNA: Hantavirus (Andes virus), LASV: Lassa virus, IAV: Influenza A virus, RSV: Respiratory syncytial virus, MARV: Marburg virus, MuV: Mumps virus, NiV: Nipah virus, RABV: Rabies virus, MeV: Measles virus, EBOV: Ebola virus. Welch's t test used to compare the difference in RNA folding of human (less enriched) cellular mRNAs versus each viral RNA (positive sense).

**F.** Comparison of Ebola minigenome vRNA generated by co-expressed, cytoplasmic T7 polymerase (no L control) and by viral polymerase L-mediated active replication inside NP marked viral condensates (+L). Confocal microscopy of fixed and stained HEK293T cell transfected with Ebola virus minigenome system, which relies on the co-expressed T7 polymerase to transcribe minigenome vRNA template from plasmid DNA. The viral polymerase L uses this T7 polymerase-generated template to replicate more vRNA and transcribe the GFP reporter-encoding mRNA. Individual channels of magnified viral condensates are shown for minigenome viral RNA stained by SABER-FISH, viral condensates marked by immunofluorescence-stained Ebola virus NP protein. In the merge overview, GFP reporter signal is pseudocolored in cyan, endogenous ODC1 mRNA signal in magenta. Scale bar: 5 μm in the merged overview image. NP-positive viral condensates are marked by yellow dashed lines.

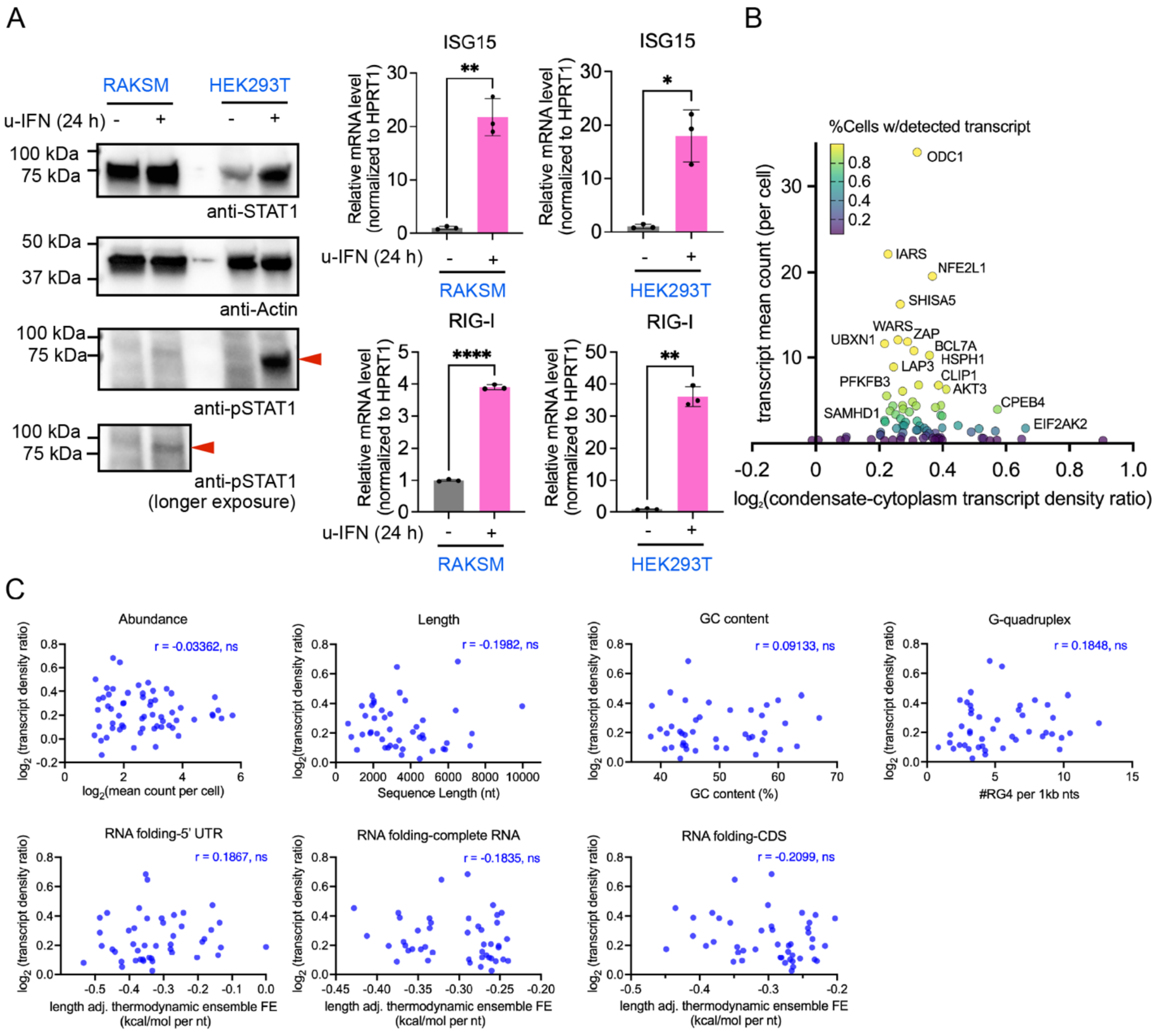


**Figure S4. Comparative MERFISH reveals altered RNA partitioning upon type 1 IFN stimulation, related to Figure 4**

**See also in Table S1**

**A.** Validation of IFN treatment-induced STAT1 phosphorylation and two representative ISGs (ISG15 and RIG-I) induction in bat kidney cells RAKSM and human kidney cells HEK 293T incubated with universal type 1 IFN (1000U/ml) for 24 hours. Whole cell lysates from equal numbers of cells in each condition were analyzed. Beta-actin used as loading control in western blotting. Arrowhead in red marks the phosphorylated STAT1 band. ISG15 and RIG-I mRNA levels were normalized to that of the house keeping gene HPRT1 mRNA level in each sample. Representative western blot and qPCR results from two experiments are shown. Unpaired t tests with Welch's correction were used to evaluate the IFN-induced ISG mRNA upregulation. *, $P<0.05$; **, $P<0.01$; **** $P<0.000$.

**B.** Distribution of log-transformed, condensate-to-cytoplasm transcript density ratios for high-confidence transcripts detected by MERFISH in the no IFN control experiment performed in parallel with the IFN-treated samples to compare IFN-mediated differential RNA partitioning. Percentage of cells with detected transcripts scaled by color map.

**C.** Correlation analysis between RNA features and condensate enrichment depicted by condensate-to-cytoplasm transcript density ratio. The Spearman r value from Spearman correlation analysis for each feature is shown. ns, not significant. Linear regression was not performed due to the lack of significant spearman correlation.

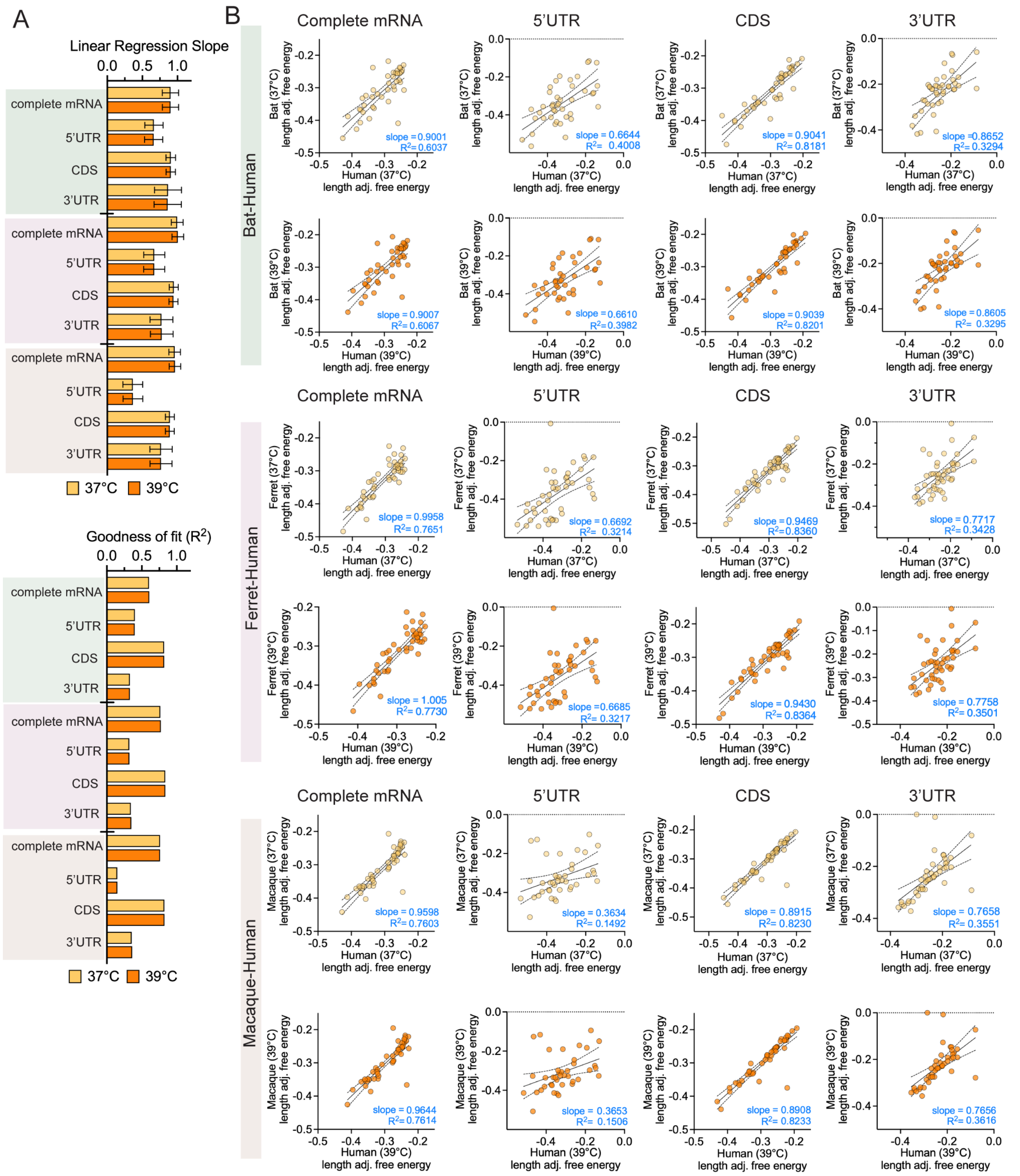

**Figure S5. Comparative RNA folding predictions for human, ferret, macaque, and fruit bat transcripts at physiological (37 °C) and fever (39 °C) temperatures, related to Figure 4**

A. Summary of linear regression metrics, including slope and $R^2$, for RNA folding of human, ferret, macaque, and fruit bat transcripts at 37 and 39°C.

B. Pairwise comparison of the degree of linear correlation between three non-human mammals (bat/ferret/macaque) and human orthologs in predicted RNA folding for 45 condensate-enriched transcripts. RNA-folding free energies of either complete RNA, 5´UTR, CDS, or 3´UTR are normalized by transcript length. OAS2 and OAS3 are expressed as a fusion gene sharing the same UTRs in bat. Physiological (37 °C) and fever (39 °C) temperatures were used for RNA-folding energy parameters and indicated by different colors. Slope and goodness of fit ($R^2$) from linear regression are shown for fitted slope with significant deviation from zero.
